# Outer membrane changes enable evolutionary escape from bacterial predation

**DOI:** 10.1101/2024.09.27.615459

**Authors:** Subham Mridha, Sarah-Luise Loose, Ove Ch. Mattmann, Saskia Isele, Rolf Kümmerli, Simona G. Huwiler

## Abstract

To combat antimicrobial resistant pathogens, natural predatory bacteria, like *Bdellovibrio bacteriovorus*, represent potential alternatives. *B. bacteriovorus* could be particularly potent as it kills a broad range of human bacterial pathogens, however, it remains unclear whether prey can evolve genetically-determined resistance against predation. Here, we show that the model bacterium *Escherichia coli* K-12 consistently evolves resistance against *B. bacteriovorus* during experimental evolution. Selection for resistance scaled positively with predation pressure and was widespread after two cycles of predator exposure. Like antibiotics, predation resistance was costly, manifesting in a trade-off between predation resistance and fitness in the absence of predators. Genetic analysis identified changes in outer membrane porin OmpF as common resistance mechanism, while a mutation in cell envelope lipopolysaccharide-modifying enzyme WaaF was rarer but also conferred predation resistance. Our study uncovers evolutionary and mechanistic aspects of prey escape from predation, generating important knowledge on predator-prey interactions and to advance sustainable treatments.

## Introduction

Bacterial resistance to conventional antibiotics is a major public health concern, that led to 1.27 million deaths in 2019^1^. This urgently calls for the discovery of novel antibiotics and alternative treatment strategies^2–4^. One promising alternative involves the use of natural bacterial predators including bacteriophages (viruses of bacteria)^5^ and predatory bacteria^3,6^, which are environmentally widespread^7^. While phage therapy is currently extensively studied^2^, much less is known about the potential of predatory bacteria, like *Bdellovibrio bacteriovorus*, that kill and invade pathogenic bacteria and are therefore considered ‘living antibiotics’^3,6^. *B. bacteriovorus* is capable of killing growing and stationary phase bacterial prey cells, and in addition irrespective of their antibiotic resistance^8–10^.

Similar to bacteriophages, *B. bacteriovorus* is neither toxic nor pathogenic to eukaryotic cells and is therefore of great promise^3,6^. However, key differences exist between the two predator types. For example, bacteriophages are typically very specific to prey strains, whereas *B. bacteriovorus* is a generalist with a broad prey range^11,12^. This makes *B. bacteriovorus* suitable to treat polymicrobial infections^11,13^. Moreover, there seem to be few defence and immunity mechanisms present among prey species to defend themselves against *B. bacteriovorus*^14,15^. This contrasts with the over hundred specific bacterial defence and immune systems reported against bacteriophages^16^. Thus, *B. bacteriovorus* might be a more effective treatment under certain circumstances, compared to traditional antibiotics and bacteriophages.

Despite these promising aspects, little is known on the selection pressures that *B. bacteriovorus* may impose on its prey and the evolutionary responses it may trigger^17–19^. Some evolutionary studies, focussing on predator-prey interactions, reported evidence for resistance evolution^17–19^ but did not explore the underlying mechanisms on a phenotypic and genotypic level. While recognition of the prey and prey invasion is likely multifactorial^20,21^, potentially hindering fast resistance formation, prey resistance has been described transient^3,22^. One study described a form of phenotypic (plastic) resistance, whereby several prey species developed a transient yet reversible stage of predation resilience^22^. Finally, several studies highlighted that secretion systems^23^ and secreted compounds such as quorum-sensing signals^24,25^, indole^26^ and cyanide^27^ may offer some protection from *B. bacterivorous* predation, while the presence of paracrystalline protein surface (S-) layers seems to prevent killing in certain bacterial species^14^.

To obtain a systematic and in-depth understanding of the potential of prey to evolve resistance to *B. bacteriovorus*, we performed an experimental prey evolution study with the model organism *E. coli* K-12 as prey and *B. bacteriovorus* HD100 as predator. Upon repeated and intermittent exposure of prey to different predator concentrations, we found strong evidence that *E. coli* K-12 predominantly evolves genetically-based predation resistance under high predatory pressure, whereas increased growth due to media adaptation and phenotypic effects played minor roles. Evolutionary escape from predation evolved repeatedly across independently evolved prey lineages. Predation resistance is costly and manifested in a negative association between the resistance level of a lineage and its growth in the absence of predation. A combination of population sequencing with high genome coverage and single-gene deletion mutant experiments revealed that mutations in the outer membrane porin OmpF, and an enzyme (WaaF) involved in lipopolysaccharide (LPS) modification are involved with resistant phenotypes. These results suggest that the prey modifies different potential points of contact with the predator^15,20^. Our study elucidates the ecological context, the evolutionary path, and the molecular mechanisms of prey resistance evolution against *B. bacteriovorus*, generating key knowledge for the development of bacterial predators as sustainable ‘living antibiotics’.

## Results

### Experimental evolution leads to increased prey growth

We experimentally evolved *E. coli* K-12 prey under no, low and high predatory pressures of *B. bacteriovorus* HD100 in eightfold replication. Experimental prey evolution was conducted for 15 alternate cycles of predation and prey recovery (Fig. 1a, Extended Data Fig. 1). After prey evolution, we subjected all evolved prey lineages to the predation pressure they experienced during experimental evolution. We found significant growth differences across the three predatory pressures (ANOVA: F_2,141 =_ 8780.2, p < 0.0001). All *E. coli* lineages evolved without predator grew significantly better than the ancestor (red dotted line), which indicates media adaptation (Fig. 1b+e). For prey lineages evolved with predators, we observed that five low-predator evolved lineages (L1-L5, Fig. 1c+f) and all high-predator evolved lineages (H1-H8, Fig. 1d+g) showed significant growth recovery relative to the ancestor subjected to predation (black dotted line). For most of these lineages (84.6%), growth recovery was partial, while only two lineages (H4, H5, 15.4%) showed complete recovery, compared to the unchallenged ancestor. Moreover, growth under predation was higher for prey lineages evolved under high than under low predatory pressure. Altogether, these results suggest that prey evolved partial resistance to predation and that higher predation pressure selected for higher levels of resistance.

**Fig. 1.**
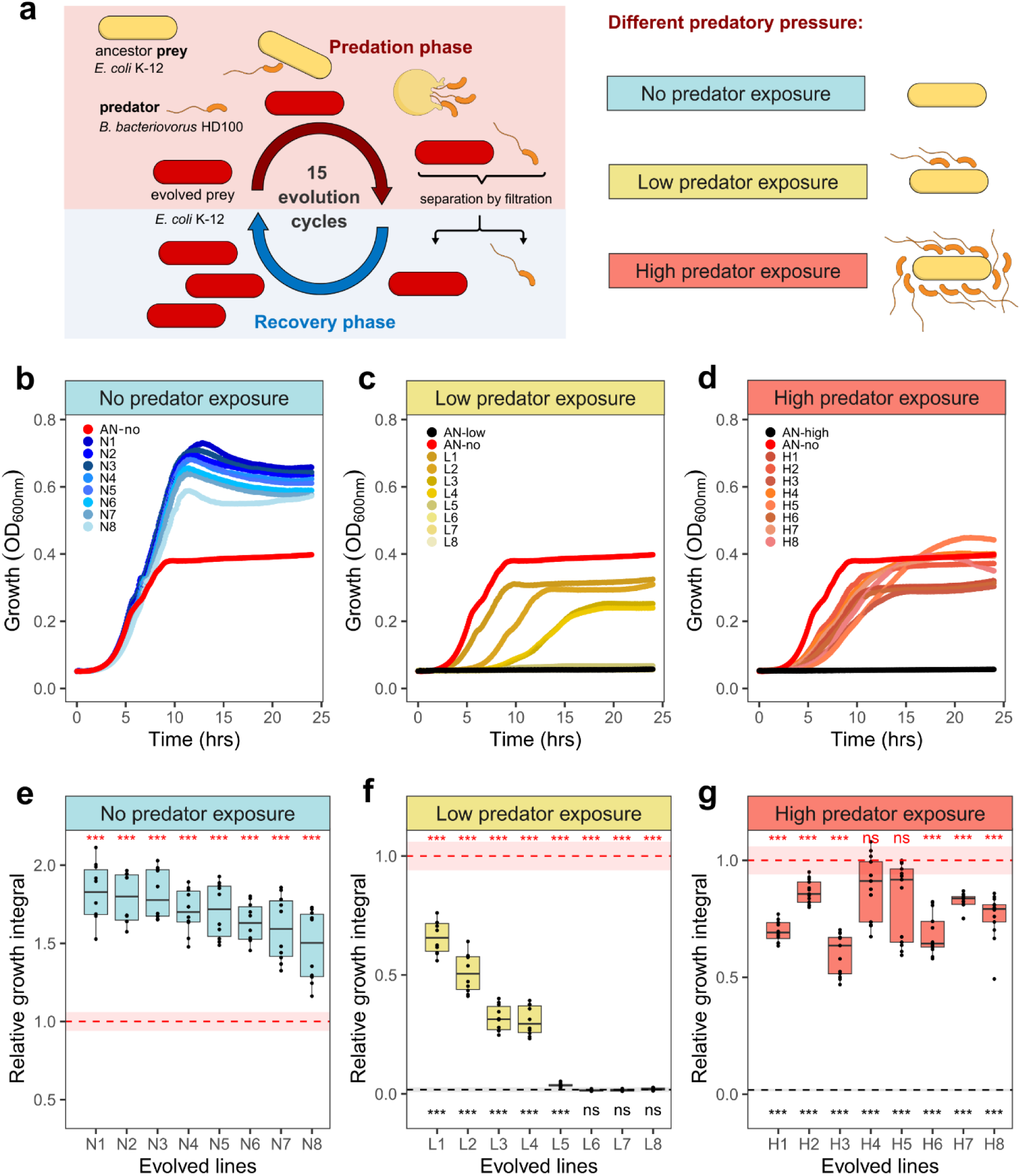
Repeated exposure of predator *B. bacteriovorus* HD100 leads to growth of prey *E. coli* K-12 under predation. **a,** Experimental evolution scheme: eight different prey lineages are exposed repeatedly to different concentration of the predator: no (N1-N8, blue), low (L1-L8, yellow) and high (H1-H8, red) predatory pressure. Each prey evolution cycle is composed of an alternating predation and recovery phase, whereby surviving prey is separated, recovered and re-exposed to predation for 15 cycles. Growth dynamics of evolved prey from 15^th^ cycle: **b-d,** growth curves measured as OD_600nm o_ver time of eight independently evolved prey lineages under no (b), low (c) and high (d) predatory pressure _when e_xposed to respective predation conditions where they evolved. They were compared with ancestor under no (AN-no) or specific predatory pressures (AN-low & AN-high)_. E_ach point of the growth curves represents average OD_600nm a_t that timepoint across replicates. **e-g,** Relative growth integral of **b-d**, defined as growth integral (area under the curve) divided by the growth integral of ancestor under no predatory pressure. Boxplots represent the median with 25^th^ and 75^th^ percentiles, and whiskers show the 1.5 interquartile range. Red dotted line and shadow indicate the mean and standard deviation range of AN-no. Black dotted lines and shaded areas indicate the mean and standard deviation range of AN under respective predation conditions. The p-value significance levels (two-sided one sample t-tests, p>0.05 = ns, p<0.05 = *, p<0.01 = **, p<0.001= ***) in comparison to AN-no is shown by red asterisks and comparison to ancestor under respective predatory pressures are shown by black asterisks. Experiments were repeated independently at least twice, N1-8 (n = 10 replicates), L1-8 (n = 10 replicates), H1-8 (n = 13 replicates), AN-no (n = 72 replicates), AN-low (n = 10 replicates) and AN-high (n = 62 replicates).

### Predation and not media adaptation drives resistance evolution

Media adaptation could be a simple explanation for the observed resistance phenotypes because faster growth would allow the prey to outpace the predator’s killing speed. If this hypothesis holds true we expect that no-predator evolved lineages, which showed highly improved growth, should be able to resist predation. We found no support for this hypothesis as all no-predator evolved lineages were still fully susceptible to predation (Fig. 2a, c). Instead, our results show that lineages L1-L5 (which evolved partial resistance under low predation) were also able to grow under high predatory pressure (Fig. 2b, d). We even observed slightly increased growth in a previously unnoticed lineage (L8). Together, these results indicate that adaptations are predation-specific and not driven by media adaptation.

**Fig. 2.**
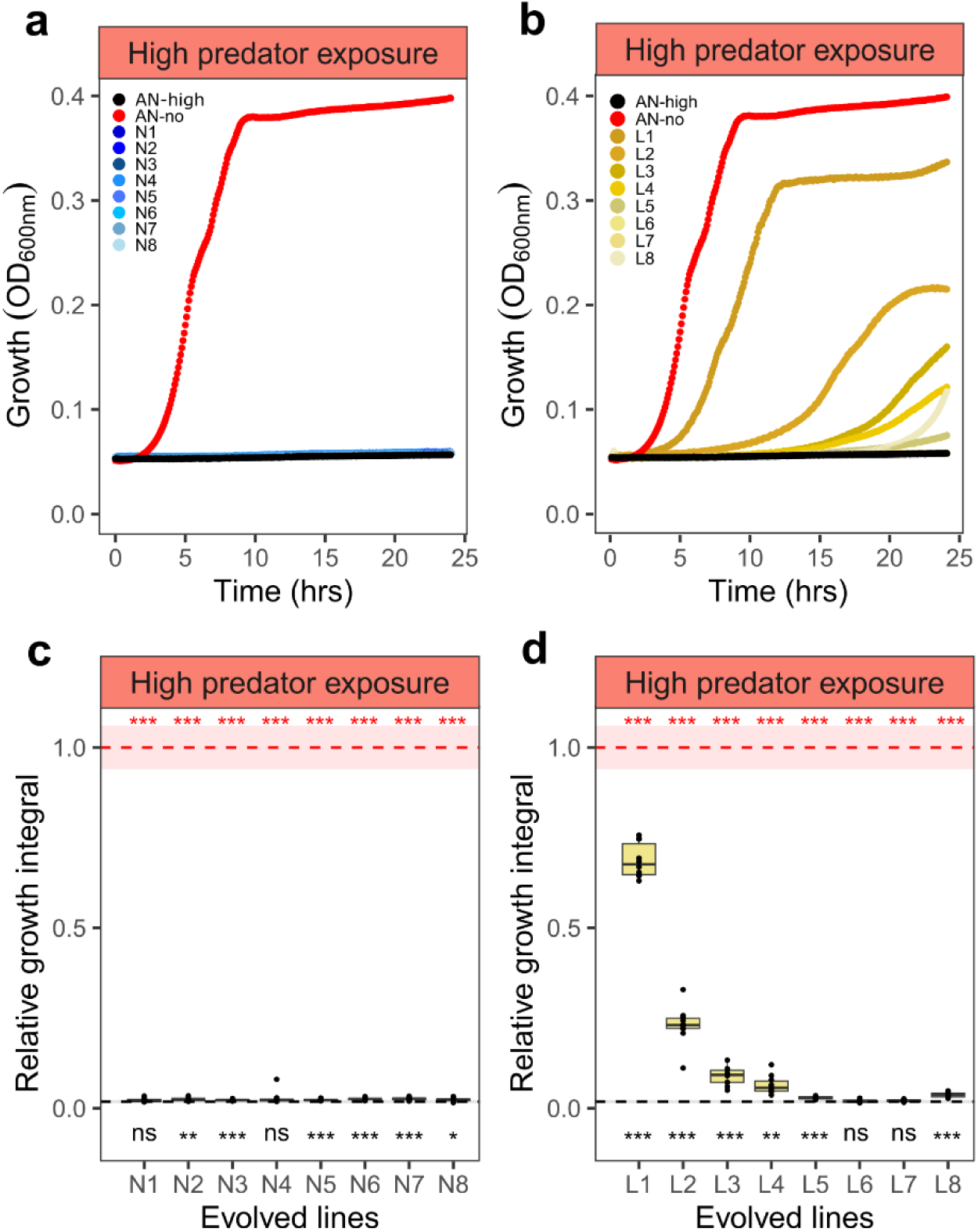
Predation resistance evolves only under predation. Growth dynamics of evolved prey from 15^th^ cycle: **a** & **b**, growth curves measured as OD_600nm o_ver time of eight independently evolved prey lineages under no (a, N1-8, blue) and low (b, L1-8, yellow) predatory pressure when exposed to high predatory pressure and compared with ancestor population under no (AN-no, red line) and high (AN-high, black line) predatory pressure_. E_ach point of the growth curves represents average OD_600nm a_t that timepoint across replicates. **c** & **d**, Relative growth integral of **a** & **b**, defined as growth integral (area under the curve) divided by the growth integral of AN-no. Boxplots represent the median with 25^th^ and 75^th^ percentiles, and whiskers show the 1.5 interquartile range. Dotted lines and shaded areas indicate the mean and standard deviation range of the ancestor under no (AN-no, red) and high (AN-high, black) predatory pressure. The p-value significance levels (two-sided one sample t-tests, p>0.05 = ns, p<0.05 = *, p<0.01 = **, p<0.001= ***) in comparison to AN-no are shown by red asterisks, and to AN-high are shown by black asterisks. Experiments were repeated independently at least twice, N1-8 (n= 10 replicates), L1-8 (n= 10 replicates), AN-no (n= 72 replicates), and AN-high (n= 62 replicates).

### Predation resistance arises early and remains persistent over time

Next, we examined the temporal pattern of predation resistance evolution. We noticed that prey optical density (OD_600nm)_ already increased at the second evolution cycle and fluctuated at a relatively high level thereafter (Extended Data Fig. 1a). To verify that resistance arose early, we repeated the growth experiments shown in Fig. 1b-g with all high-predator lineages (H1-H8) from the 2^nd^ and 5^th^ evolution cycle (Extended Data Fig. 1 and 2). These analyses confirmed that predation-resistant phenotypes emerged early on during experimental evolution as all lineages could readily grow under high predation pressure at evolution cycle 2, with barely any growth differences existing compared to later evolution cycles 5 and 15 (Extended Data Fig. 1 and 2). Hence, predation resistance evolved early and once emerged within a prey lineage it remained persistent over generations.

### Trade-off between predation resistance and prey growth in the absence of predation

Antibiotics resistance is often associated with fitness costs in the absence of antibiotics^28,29^. Here, we tested whether a similar trade-off occurs for *E. coli* in the context of predation resistance. To investigate this, we grew all evolved lineages without predators and calculated their growth rate relative to the ancestor. Relative growth rates differed significantly across predatory pressures (ANOVA: F_2,141 =_ 460.42, p < 0.0001), and were highest in no-predator evolved *E. coli* lineages (Fig. 3a, d), followed by low-predator evolved lineages with growth rates similar to the ancestor (Fig. 3b, e) and high-predator evolved lineages with generally lower growth rate than the ancestor (Fig. 3c, f). When contrasting these growth rates against relative prey resistance (quantified as relative growth integral under high predatory pressure), we found a significant negative correlation between predation resistance and prey fitness, indicating the predicted trade-off (Fig. 3g). Thus, our findings corroborate that *E. coli* resistance mechanisms are costly in the absence of *B. bacteriovorus* predation.

**Fig. 3.**
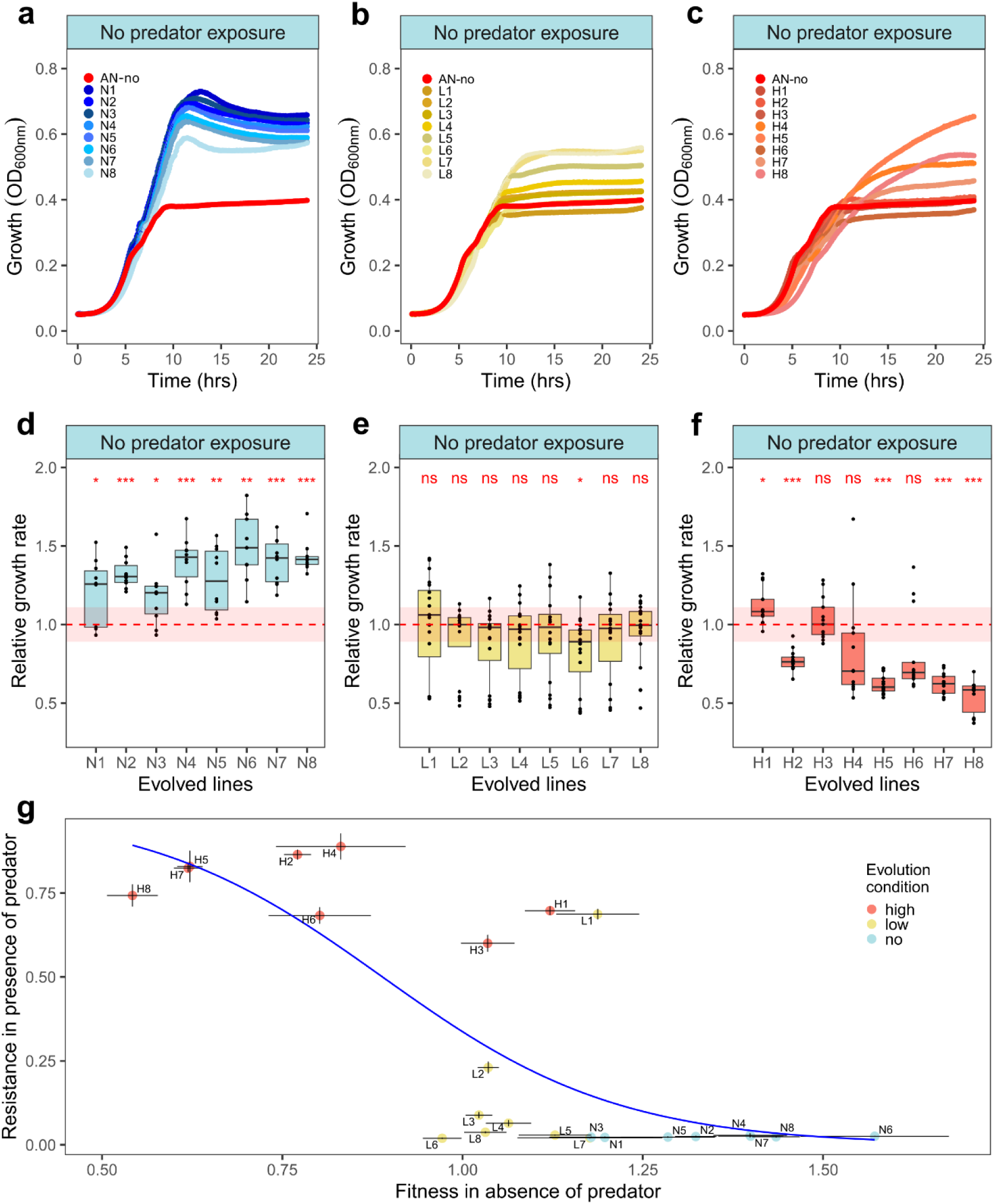
Prey resistance to bacterial predator is generally associated with negative fitness consequences. Growth dynamics of evolved prey from 15th cycle: **a-c** growth curves measured as OD600nm over time, of prey lineages evolved under no (**a,** blue), low (**b,** yellow) and high (**c,** salmon) predatory pressure, when exposed to no predatory pressure. Each point of the growth curve represents the average OD600nm at that timepoint across replicates. Panel **a** is identical with Fig. 1a for a complete overview. **d-f,** Relative growth rates of no (**d**), low (**e**), and high (**f**) predatory pressure evolved lines, when exposed to no predatory pressure. Relative growth rate is defined as maximum growth rate divided by the maximum growth rate of the ancestor under no predatory pressure (AN-no). Boxplots represent the median with 25th and 75th percentiles, and whiskers show the 1.5 interquartile range. Red dotted lines and shaded areas indicate the mean and standard deviation range of ancestor under no predatory pressure (AN-no). The p-value significance levels (two-sided one sample t-tests, p>0.05 = ns, p<0.05 = *, p<0.01 = **, p<0.001 = ***) in comparison to AN-no are shown by red asterisks. Experiments were repeated independently at least twice, N1-8 (n= 10 replicates), L1-8 (n= 20 replicates), H1-8 (n= 13 replicates), AN-no (n= 72 replicates). Three outliers, one each from N6, L4 and L7 were excluded from panels **d & e** for better visualisation but were included in the statistical analysis. **g,** Trade-off between predation resistance and prey fitness. Fitness, quantified as mean relative growth rate under no predatory pressure (**d-f**) versus predator resistance, quantified as mean relative growth integral under high predatory pressure (Fig. 1g & 2c-d). N1-8, L1-8, and H1-8 denotes prey lineages that evolved under no (blue), low (yellow) and high (salmon) predatory pressure respectively. A logistic regression function was fitted as a blue line to correlate the two variables (resistance versus fitness, Wald z-statistics = -2.336, p-value = 0.0195). Black error bars represent the standard error of mean for the respective variables.

### High consistency in resistance levels among single clones

Based on the detected trade-off between predation resistance and population prey growth, we wondered whether there is heterogeneity in predation resistance among clones within prey populations. Consequently, we analysed the predation resistance profile of six randomly isolated clones from each of the eight high-predator lineages (15^th^ cycle, 48 clones) and found that all clones showed consistently high predation resistance. This shows that clones did not separate into susceptible and resistant phenotypes (Extended Data Fig. 4 and 5) and thus did not segregate along a trade-off line within populations. This result further indicates that the mechanisms conferring resistance to predation likely operate at the individual and not at the group level of the prey. Such group-level mechanisms were proposed to operate via the secretion of small soluble substances (quorum-sensing signals, indole, cyanide)^24–27^ and are expected to protect not only the evolved producers but also susceptible wildtype cells.

### Mutation frequency in the prey genome peaks under high predatory pressure

To uncover the genetic basis of putative resistance mechanisms, we sequenced all 24 evolved *E. coli* lineages (N1-8: no predation, L1-8: low predation, H1-8: high predation) after the 15^th^ evolution cycle along with the original ancestor with high sequencing depth. The Illumina sequencing reads were mapped to the reference genome of *E. coli* K-12 MG1655 and multi-sample variant calling with a frequency cutoff of 5% was performed. We focused on *E. coli* genes that had mutations in evolved lineages but not in the ancestor (Extended Data Fig. 6). We excluded mutated rRNA and transposase genes because they did not correlate with any of the three evolution conditions (Extended Data Fig. 7). Using these filters, we uncovered 43 mutated genes of which 17 occurred in N-lineages, 17 in L-lineages and 30 in H-lineages (Fig. 4a). A total of 26 genes mutated exclusively under predatory pressure (Fig. 4a), and thus reflect the top candidates potentially explaining resistance phenotypes. H-lineages, had more exclusive gene mutations than L-lineages (18 vs. 5, Fig. 4b). Moreover, there were fewer high-frequency mutations (>70%) among L-lineages than among H-lineages (Fig. 4b), which is likely explained by the higher selection pressure.

**Fig. 4.**
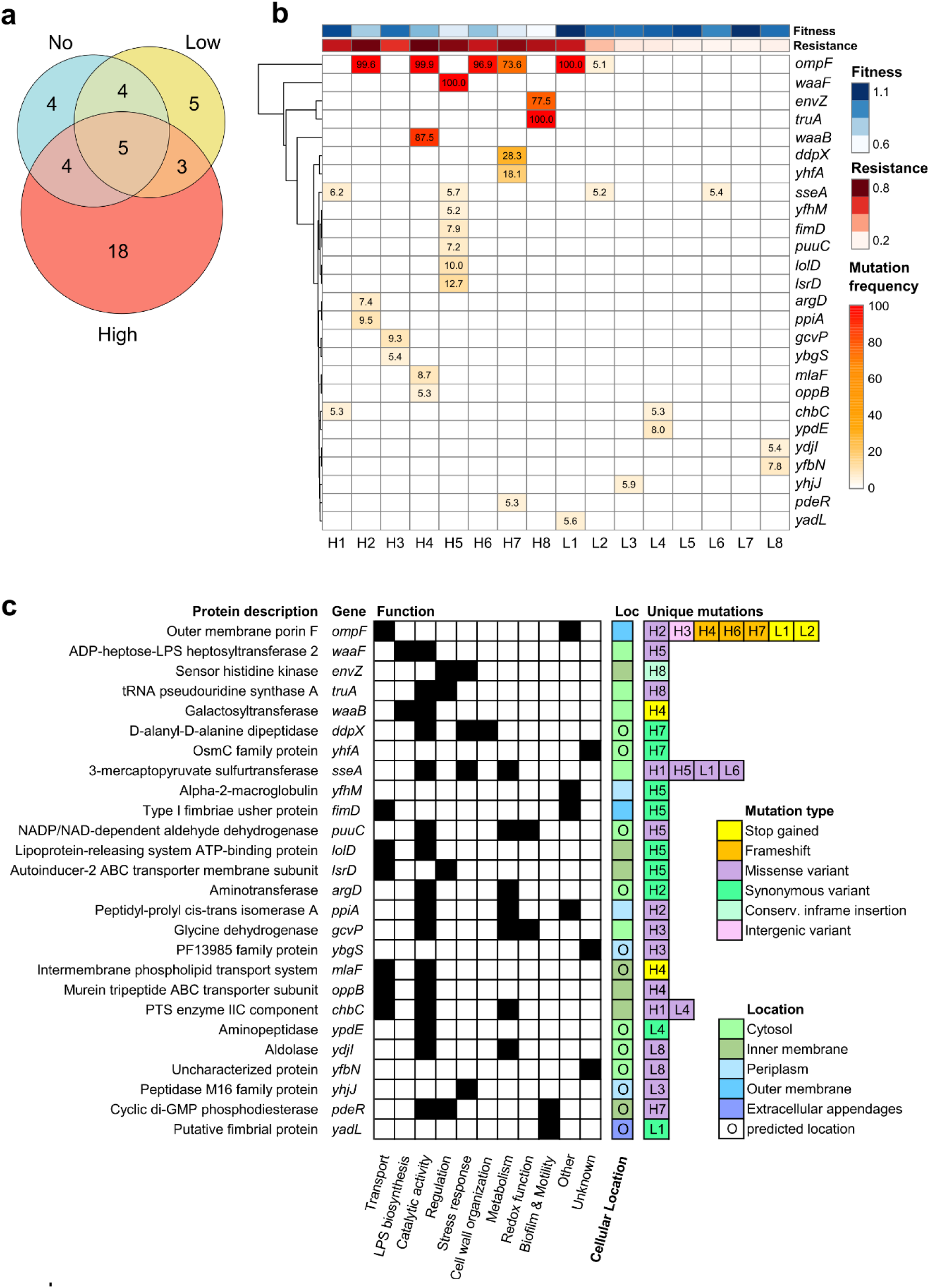
Gene mutational patterns in prey populations evolved under repeated predation pressure. Genes mutated in evolved *E. coli* K-12 lineages after the 15^th^ evolution cycle in comparison to the ancestor. **a,** Venn diagram showing the number of mutated *E. coli* genes that reached frequencies higher than 5% within a lineage, grouped according to no, low and high predatory pressure. Not included in this Venn diagram are mutated rRNA and transposase genes, which are listed separately (Extended Data Fig. 7). **b**, Hierarchically clustered heatmap of mutation frequency differences relative to the ancestor of all the genes that exclusively mutated under high (H1-8) and/or low (L1-8) predatory pressure. Heatmaps on top of the panel show fitness (blue) measured as average relative growth rate of lineages under no predatory pressure, and resistance (red) measured as average relative growth integral of lineages under high predatory pressure, according to the data in Fig. 3g. **c**, Overview of mutated genes and mutation types together with a description of the encoded proteins, their functions and cellular location (Loc) for all the genes that exclusively mutated under high and/or low predatory pressure, according to **b**.

### Outer membrane porin OmpF mutations repeatedly arose and confer resistance to predation

Next, we conducted a cluster analysis (Fig. 4b) and analyzed the functions of the mutated gene products including mutation types and (predicted) amino acid changes (Fig. 4c, Extended Data Table 1). These analyses yielded *ompF*, encoding the outer membrane porin F, as the most frequently mutated gene. Four H-lineages (H2, H4, H6 & H7) and two L-lineages (L1, L2) had mutations in this gene. Moreover, lineage H3 had a point mutation in the upstream region of *ompF* (Extended Data Fig. 8) and lineage H8 had an inframe insertion in *envZ* (Extended Data Table 1), which encodes a sensor histidine kinase involved in OmpF regulation^30^. Mutation frequencies were high (73.6%-100%) in all lineages except in L2. The sequencing of six individual clones per H-lineage showed that mutations in *ompF* and *envZ* were present in all clones (Extended Data Fig. 9). Overall, there were no major differences in mutational patterns between clonal and population level sequencing (Extended Data Fig. 9), reinforcing the view of selective sweeps of a few beneficial mutations through populations. Mutation types in *ompF* include missense variants (H2), frameshifts (H4, H6, H7) and direct stop gains (L1, L2) (Fig. 4b, c, Extended Data Fig. 8, Extended Data Table 1), indicating that most observed mutations probably lead to OmpF function loss.

To directly test whether the loss of OmpF (a known attachment side for phages and known to be critical for effective *B. bacteriovorus* predation^15^) confers resistance to predation, we generated a clean, single deletion Δ*ompF* mutant based on the *E. coli* KEIO library (details in methods, Supplementary Online material Table SOnline1). First, we confirmed that *E. coli* KEIO reference strain (BW25113) was fully susceptible to *B. bacteriovorus* and grew well in the absence of predation (Fig. 5a,b), thus showing similar responses as the *E. coli* K-12 ancestor (Fig. 1b,d and 2a,b). We then subjected the Δ*ompF* mutant to no and high predation pressures and tracked its growth. We found this mutant to be completely resistant to predation, showing full growth recovery under high predator exposure (Fig. 5a,b). We repeated the above experiments with a Δ*envZ* clean deletion mutant and found that this mutant was still susceptible to predation, in agreement with Mun et al.^15^ (Fig. 5a, b). This is unsurprising because the experimentally evolved *envZ* mutant involved an inframe insertion [insertion of Lys-Thr-Trp-Leu between Leu135 and Lys136] and not a deletion. We suspect that the insertional mutation in our experiment might upregulate OmpR, resulting in lower levels of OmpF. We thus conclude that functional changes in EnvZ, but not loss-of-function, could confer predation resistance. In summary, our results indicate that predation resistance is primarily driven by OmpF loss-of-function mutations.

**Fig. 5.**
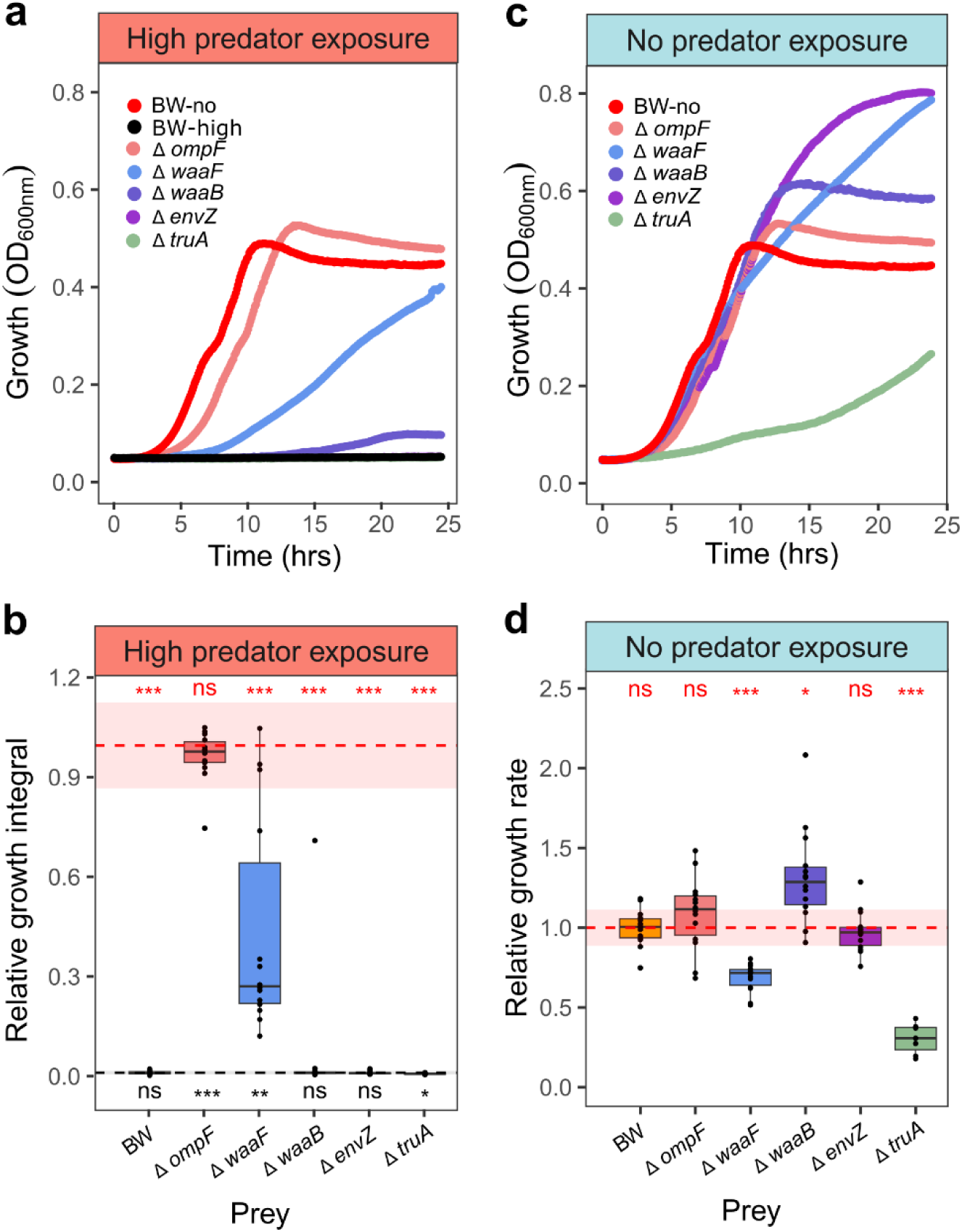
OmpF deletion in *E. coli* prey results in high predation resistance. Different *E. coli* clean-deletion mutant strains (Δ*ompF,* Δ*waaF,* Δ*waaB,* Δ*envZ,* Δ*truA*) were generated based on the KEIO library. *E. coli* BW25113 (BW) is the KEIO reference strain. **a+c**, Growth curves _measured as O_D_600nm o_ver time of the different *E. coli* deletion mutants exposed to either high predatory pressure (**a**) or no predators (**c**). Each point of the growth curve represents the average OD_600nm p_er timepoint across replicates. Experiments were independently repeated at least twice. Total replicate numbers for the various strains: Δ*ompF* (n= 14), Δ*waaF* (n= 14), Δ*waaB* (n= 14), Δ*envZ* (n= 14), Δ*truA* (n= 8), BW without predatory pressure (BW-no) (n= 14), and BW with high predatory pressure (BW-high) (n= 15). **b,** Relative growth integrals from **a**. Relative growth integrals are calculated as growth integral (area under the curve) of the clean-deletion mutant with predatory pressure divided by the growth integral of BW-no. **d**, Relative growth rates from **c**. Relative growth rates are calculated as the maximum growth rate of the clean-deletion mutant divided by the maximum growth rate of BW-no. Boxplots represent the median with 25^th^ and 75^th^ percentiles, and whiskers show the 1.5 interquartile range. Red and black dotted lines and shaded areas indicate the mean and standard deviation range of BW-no and BW-high respectively. The p-value significance levels (two-sided one sample t-tests, p>0.05 = ns, p<0.05 = *, p<0.01 = **, p<0.001= ***) in comparison to BW-no (red asterisks), and to BW-high (black asterisks).

### Mutations in *truA* cause costly prey morphology change

In lineage H8, a high frequency mutation occurred in *truA* (100%, missense: Val200Ala) in addition to the above-described *envZ* mutation (77.5%). The *truA* gene encodes the tRNA pseudouridine synthase A important for protein synthesis^31^. Phase-contrast microscopy revealed that cells from lineage H8 no longer separated properly after cell division resulting in cells being thinner and longer (Extended Data Fig. 3a-c). Experiments with a Δ*truA* clean deletion mutant confirmed division abnormalities (Extended Data Fig. 3), matching previously reported results on the effect of *truA* disruption^32^. No such abnormalities in cell morphology occurred in the other H- and L-lineages (Extended Data Fig. 3, Fig. SOnline1). However, deletion of *truA* did not result in predation resistance but was rather associated with severe fitness costs in the absence of predation (Fig. 5). We hypothesise that the resistance phenotype observed in the evolved *truA* mutant is masked in the clean deletion mutant due the detrimental effects on cell division (Fig. 5). Further, our mutation frequency data suggest that the missense mutation in *truA* (100%) happened first and was associated with a drop in final OD_600nm a_fter the recovery phase of early evolution cycles 1-5. Growth only recovered in later cycles, likely due to the additional mutation in *envZ* (77.5%), probably partly compensating the fitness drop (Extended Data Fig. 1b). Altogether, our results indicate that resistance to predation may occur through cell morphology changes. However, such events might be rare due to the severe fitness costs typically associated with changes in fundamental cell physiological processes.

### Mutation in LPS synthesis gene *waaF* confers partial resistance to predation

In lineage H5, a high frequency mutation occurred in *waaF* (100%, missense: Ala255Glu), encoding an heptosyltransferase involved in the biosynthesis of the lipopolysaccharide (LPS) inner core^33^. We hypothesized that modification or abolishment of outer membrane structures could represent a mechanism through which *E coli* can become resistant to predation. In support of our hypothesis, we found that a Δ*waaF* clean deletion mutant showed partial and significant resistance to predation (Fig. 5a,b). The same hypothesis could also hold true for the high frequency mutation observed in *waaB* (lineage H4, 87.5%, stop codon gained) encoding a galactosyltransferase involved in LPS biosynthesis. However, contrary to our expectation we did not find Δ*waaB* (clean deletion mutation) to be resistant to predation (Fig. 5a,b). Instead, it showed significant growth improvements in the absence of predation (Fig. 5c,d). Because the mutation in *waaB* occurred together with an *ompF* mutation in lineage H4 we conclude that WaaB loss-of-function does not confer predation resistance but is beneficial in an already resistant lineage, showing the highest growth recovery under predation among all H-lineages (Fig. 1g).

## Discussion

Historical records on antibiotics tell us that bacterial pathogens have evolved resistance to every single drug used in clinics. Nowadays, it is therefore imperative to understand putative mechanisms and paths to resistance already during the development of new antibacterials^34,35^. In our study, we performed this task for the predatory bacterium *B. bacteriovorus*, which is regarded as a potential ‘living antibiotic’ preying on human pathogens. We found that the model prey *E. coli* K-12 MG1655 consistently evolved resistance against predation through two main mechanisms: change in the outer membrane porin OmpF (several independent events), and change in WaaF (one event), an enzyme involved in outer membrane LPS biosynthesis. These mechanisms suggest that selection acts on cell-membrane associated traits that form the first line of contact between prey and predators. The uncovered resistance mechanisms are similar to the ones observed to evolve against bacteriophages^36,37^ (loss/alteration of host receptors^38,39^) and against membrane-targeting polymyxin antibiotics (LPS modifications)^40^.

OmpF enables *E. coli* to import and export small molecules via passive diffusion. Osmolarity of the environment likely effects prey susceptibility towards *B. bacteriovous*. It is known that *E. coli* expresses high levels of OmpF under neutral pH, whereas high osmolarity leads to its downregulation^41^. OmpF serves as an entry point for bacterial toxins (colicin) and antibiotics^42–44^ while simultaneously serving as docking site for certain bacteriophages^38,39^. However, its role in the interaction with *B. bacteriovorus* is not yet resolved, apart from the fact that the deletion of OmpF slows predation rate^15^ and that mutations in this gene are under selection in response to predation in our study. We can thus only speculate about how loss or impairment of OmpF could foster predation resistance, and we propose three scenarios. Firstly, change of OmpF could prevent predators to use the porin as a docking site for attachment^15^, similar to phage invasion^38,39^. Secondly, OmpF might serve as an ‘overpressure valve’ equilibrating the osmotic pressure building up when *B. bacteriovorus* enters the prey periplasm. Loss of OmpF or a reduction in its small molecule transport rate might compromise this function and presumably slow down predator entry. Thirdly, loss of OmpF could reduce the entry of potential, hypothetical small molecules secreted by the predator. Such molecules could manipulate the prey during attachment and recent evidence indicates that *B. bacteriovorus* can kill the prey even without subsequent entry^45^. Altogether, our results combined with previous reports indicate that OmpF is a mutational hotspot in the context of resistance evolution against antibiotics, bacteriophages, and predatory *B. bacteriovorus*.

Important to note is that the relevance of OmpF for successful *B. bacteriovorus* predation seems to depend on the prey species^15^. For example, deletion of OmpF in *Salmonella enterica* did not reduce prey killing^15^. Instead, it was reported that the LPS core has a major influence on attachment^46,47^. This finding combined with our results (showing that mutations in LPS-modifying enzymes WaaF and WaaB were under selection) strongly support the view that LPS structure takes on an important role in the interactions with predators. Work with bacteriophages revealed two opposite roles for LPS (particularly the long O-antigens): depending on their specific structure they may act as primary receptors for attachment or block attachment to the outer membrane^38,39^. Several lines of evidence suggest that the dual role also applies to interactions with *B. bacteriovorus*^20,46,48^. First, a phage-tail fiber protein homolog of *B. bacteriovorus* was indeed shown to bind to multiple bacterial surface glycans/LPS in multiple prey species (*E. coli* K-235, *Serratia marcescens* and *Proteus mirabilis*)^20^. Second, attachment to prey is slowed down when predators are confronted with complex O-antigens as opposed to shorter LPS only featuring the core oligosaccharides in several species (*S. enterica*, *E. coli*, *Vibrio cholerae*)^46^ ^48^. In sum, our results indicate that evolutionary change in LPS structure can modulate prey-predator interactions as observed for bacteriophage-bacteria interactions.

At first sight, our results on repeated resistance evolution seem to dampen the hope for predatory bacteria to become sustainable antibacterial treatments. Nonetheless, our experiments also revealed good news. Particularly, the strong fitness trade-off between resistance to predation and growth in the absence of predation highlights that genetically manifested predation resistance is costly, compromising cellular functionality. This phenomenon could limit the spread of resistant prey as they are weak competitors and probably cleared away more efficiently by macrophages. There are concepts to leverage such competitive constraints to increase treatment sustainability^28,49^. A key condition for leveraging on competitive constraints is that the pathogen is eliminated before compensatory mutations arise. We observed two putative compensatory mutations in our experiments (mutation in *waaB* in the background of an *ompF* mutant; mutation in *envZ* in the background of a *truA* mutant). Clearly, more research is required to understand the propensity of resistant prey mutants (with and without compensatory mutations) to spread. Moreover, it will be important to test whether similar evolutionary patterns (type of mutants, compensatory mutants and speed of resistance development) are detected when using predatory *B. bacteriovorus* against pathogenic bacteria and clinical isolates as prey to assess the sustainability of this ‘living antibiotic’.

To conclude, we like to emphasize two important points regarding prey resistance against bacterial predation by *B. bacteriovorus*. First, resistance mechanisms do not need to offer complete protection to escape predation. Instead, it suffices to alter the balance between prey growth and killing rate. In our experiment, we detected a decrease in killing rate (by mutations in *ompF* and *waaF*), but no increase in prey growth rate. Second, while we focused on predation resistance through mutation and selection processes, previous work has reported ‘plastic’ resistance phenotypes, occurring after short timescale exposures, offering transiently protection against *B. bacteriovorus*^22^. Such plastic effects also surface in response to colistin exposure (i.e. change in membrane polarity) and might well have occurred in our experiments (e.g., in resistant prey lineages exposed to low predatory pressure with few low frequency genetic mutations). A key task for future studies is to integrate all these aspects and to understand the dynamics of prey resistance and predation escape under situations that are closer to clinical settings – for example using clinical pathogen isolates, assessing treatment efficacy in spatially structured human cell cultures, and allowing predators to co-evolve with the prey. Only with such integrative knowledge, we can ensure the long-term sustainability of *B. bacteriovorus* as a ‘living antibiotic’, efficient in targeting multi-drug resistant pathogens, in clinically relevant contexts.

## Supporting information

Online Statistics Table

## Acknowledgments

The authors would like to thank Dr. Timothy Sykes and Dr. Weihong Qi from Functional Genomics Center Zurich (FGCZ) for DNA library preparation, Illumina sequencing and initial data analysis by multi-sample variant calling. Microscopy images were obtained with a Leica Thunder microscope maintained by the Center for Microscopy and Image Analysis at University of Zurich. We would like to thank Prof. R. E. Sockett (University of Nottingham, UK) for providing *B. bacteriovorus* HD100^T^ and helpful input on our manuscript. Further, we would like to thank Prof. Alex Hall and Dr. Daniel Angst (both ETH Zürich, CH) for the *E. coli* strains from the KEIO collection and pCP20. This study was enabled by FAN (Research Talent Development Fund of the University of Zurich Alumni), the Ernst Göhner Stiftung, as well as the Kurt und Senta Herrmann-Stiftung (all to S.M. and S.G.H.), all of which contributed towards research costs and salary of S.M.. R.K. was funded by a project grant from the Swiss National Science Foundation (no. 310030_212266), S.G.H. was funded by Swiss National Science Foundation Ambizione Fellowship PZ00P3_193401. A CC-BY-NC-ND copyright license is applied to this manuscript.

## Author contributions

S.M. and S.G.H. jointly conceived the project and obtained funding with input from R.K.. S.M. and O.Ch.M. conducted the prey evolution experiment and isolated genomic DNA from prey populations for sequencing of ancestor and evolved prey lineages. S.M. and O.Ch.M. isolated single clones of evolved prey lineages and sent them for sequencing. S.M. and O.Ch.M. conducted predation assays with prey of ancestor and evolved lineage populations at the end timepoint (15^th^ evolution cycle). S.M. made predation assays of evolved prey lineage populations at 2^nd^ and 5^th^ evolution cycles as well as evolved single clones (15^th^ evolution cycle). S.L.L. performed predation assays of prey deletion strains with help of S.I.. Data analysis and evaluation of predation assays was done by O.Ch.M., S.L.L., and S.M. (lead). Genome sequence analysis was performed by O.Ch.M. (population), S.G.H. (population, gene mutations), S.L.L. (single clones, OmpF population mutations) and S.M. (population, single clones, all final heatmaps, lead). S.L.L. generated clean deletion strains Δ*ompF,* Δ*waaF,* Δ*envZ,* and Δ*truA.* S.I. and S.G.H. helped S.L.L. to generate Δ*waaB*. S.L.L. took microscopy images of prey ancestor and all evolved prey lineage populations (15^th^ evolution cycle), as well as *E.coli* BW25113, Δ*envZ,* and Δ*truA*, with assistance of S.G.H.. S.L.L. performed image analysis with statistical input by S.M.. S.G.H. advised on predatory bacterial culturing, lab experimentation, prey sequencing, and molecular genetics aspects. R.K. advised on evolutionary and statistical aspects. S.M., S.G.H. and R.K. wrote the paper with contributions from the other authors. All authors approved of the final version before submission.

## Additional information

**Extended data** is available for this paper at https://doi.org/XXX.

**Supplementary information** The online version contains supplementary material available at https://doi.org/XXX.

**Requests for materials** should be addressed to Simona G. Huwiler.

## Methods

### Growth of bacterial strains

In this study, *Escherichia coli* K-12 MG1655 (DSM 18039) as prey and *Bdellovibrio bacteriovorus* HD100^T^ as predator were used, unless indicated otherwise. *B. bacteriovorus* HD100^T^ was obtained from the laboratory of Prof. R. E. Sockett at University of Nottingham, UK. *B. bacteriovorus* HD100 was cultured and maintained on Ca/HEPES buffer (25 mM HEPES, 2 mM CaCl_2,_ pH 7.6) with late-log *E. coli* K-12 MG1655 at 29°C for generally 24 h at 200 rpm shaking^1^. General revival of *B. bacteriovorus* and determination of plaque forming units (PFU) for enumeration was done according to protocols by Lambert & Sockett^1^ and is described in more detail at the end of the Additional Online materials.

All *E. coli* strains used are listed in Additional Online Materials Table SOnline1. To validate the genetic effect on predation resistance, markerless single deletion strains Δ*envZ,* Δ*ompF,* Δ*truA,* Δ*waaB,* and Δ*waaF* were generated based on *E.coli* KEIO library strains^2^ containing an inserted kanamycin resistance cassette (following section). The different *E. coli* strains were generally grown in YT both (Yeast tryptone, 5 g l^-1^ NaCl, 5 g l^-1^ Bacto yeast extract, 8 g l^-1^ tryptone, pH 7.5) at 37°C with 200 rpm shaking for generally 16 h overnight, and additional appropriate antibiotics as indicated. For ancestor prey, *E. coli* K-12 MG1655 was retrieved from 20% glycerol stocks at -80°C and were streaked on a YT agar plate (YT broth with 10 g l^-1^ agar) and incubated at 37°C for 24 h to obtain single colonies. The overnight cultures from ancestor prey single colonies were used for 24-hr lysates (growth of predator), revival and enumeration of the predator in our experiments.

### Generation of markerless deletion strains of *E. coli* KEIO library

Markerless single deletion mutants in *E. coli* (Δ*envZ,* Δ*ompF,* Δ*truA,* Δ*waaB,* and Δ*waaF*) were constructed of the respective strain from the KEIO collection (based on *E. coli* BW25113)^2^. To achieve this, the kanamycin resistance cassette (present in the KEIO collection to disrupt the individual genes) was removed to enable predation assays with kanamycin-sensitive *B. bacteriovorus* HD100. The kanamycin cassette was excised via pCP20^3^, expressing an FLP-FRT recombinase. To generate electrocompetent cells, and excise the kanamycin cassette, several protocols from the Barrick lab were adapted^4,5^. To prepare electrocompetent cells the desired *E. coli* strain in YT broth with 30 μg/ml kanamycin was incubated for 16 h at 37°C and 200 rpm. The stationary phase culture was used to inoculate at least 10 ml of YT broth with 30 μg/ml kanamycin at optical density with absorbance at 600nm (OD_600nm)_ of 0.05 and grown at 37°C and 200 rpm. At an OD_600nm o_f ∼ 0.6 the culture was centrifuged for 5 minutes at 6000 x *g* and 4°C, and the supernatant was discarded. The resulting pellet was washed 4 times with 10% ice-cold glycerol of the same volume as the initial inoculation volume. Finally, the pellet was resuspended in 10% glycerol at 100x concentration, and 50 μl aliquots stored at -80°C. These electrocompetent *E. coli* cells were thawed on ice and maximally 100 ng of pCP20 added. After gentle mixing, the sample was transferred into a cooled 0.2-cm cuvette (Bio-Rad), and electroporated at 2.5 kV with a MicroPulser electroporator (Bio-Rad). Then, the sample was resuspended in 500 μl warm SOC medium and incubated for 30 min at 30°C and 120 rpm. The cells were plated on YT agar plates with 100 μg/ml ampicillin and grown at 29°C for at least 16 h. To select for loss of heat-sensitive plasmid pCP20 after recombination, a single colony from a YT agar plate with 100 μg/ml ampicillin was picked to inoculate 5 ml of YT broth, and grown for 16 h at 43°C, shaking at 200 rpm. 50 μl of a 10^-6^ dilution of this stationary phase culture was plated on YT agar without antibiotics for single clones. To screen for successful recombination and loss of plasmid, six colonies were selected and streaked out on YT agar with 30 μg/ml kanamycin, YT agar with 100 μg/ml ampicillin, and YT agar without antibiotics in this order. The YT plates without and with kanamycin were incubated at 37°C, the YT plates with ampicillin at 29°C for at least 16 h. To screen for successful excision of the kanamycin cassette, recombinants with sensitivity to both antibiotics were grown in 5 ml of YT broth, incubated for 16 h at 200 rpm and 37°C. Of these cultures, 20% glycerol stocks were made and stored at -80°C. To control correct excision of the kanamycin cassette, PCR was used to assess correct product size (Primers in Additional Online Materials Table SOnline 2). Additionally, correct scar regions were confirmed by sanger sequencing (Microsynth AG, CH).

### Experimental prey evolution

In the experimental prey evolution, *E. coli* K-12 MG1655 was used as ancestral prey strain and subjected to three different predatory pressures by *B. bacteriovorus* HD100 involving 15 alternate cycles of predation and recovery phase (Fig. 1a). The detailed protocol is described at the end of the Supplementary Online Materials. The predation phase occurred in rich medium (containing ¾ YT, 1/4 25 mM HEPES and 2 mM CaCl_2)_ allowing growth of *E. coli* K-12 as well as predation of *B. bacteriovorus* at 200 rpm and 29°C. *E. coli* K-12 prey were subjected to three different predatory conditions: No predatory, low predatory pressure (multiplicity of infection [MOI], ratio of predator (PFU/ml) vs prey cells (CFU/ml), was ∼ 2.5) and high predatory pressure (MOI ∼10). For each condition eight independent prey lineages were evolved from the *E. coli* K-12 ancestor. After the 24-hr predation phase surviving *E. coli* were separated from *B. bacteriovorus* by filtration to avoid co-evolution. In the subsequent 16-hr recovery phase retrieved *E. coli* from filtration were diluted ∼1:100 and grown in YT at 200 rpm and 37°C. These overnight cultures were diluted back to an overall initial OD_600nm o_f 0.003 for the next predation phase and evolution cycle.

### Isolation of prey single clones

Prior to the experiment we made overnight cultures of evolved lineages under high predatory pressure (15^th^ evolution cycle) stored at -80°C, by extracting cell material (containing evolved population) from cryo stocks and inoculating 50 ml YT medium in 250 ml Erlenmeyer flasks, incubated at 37°C, 200 rpm for 16 h. We harvested prey cells from 10ml of these overnight cultures by centrifugation at 5311 x *g* for 3 min and subsequently washed them in 10 ml of Ca-HEPES. Then prey cells were resuspended in 6 ml of Ca-HEPES and adjusted to OD_600nm =_ 3.0, then we diluted the cultures to 10^-6^ and plated 100 μl on YT agar plates and incubated them at 37°C for 24 h. We randomly selected 6 single colonies from each evolved lineage and overnight cultures were made with these single colonies in 10 ml YT medium in 50 ml Falcon tubes incubated at 37°C, 200 rpm for 16 h. From this culture we prepared 20% glycerol stocks and stored them at -80°C. Prior to experiments, we made overnight cultures of the single colonies from cryo stocks, alongside their respective evolved lineage population and ancestor in 50 ml YT medium in 250 ml Erlenmeyer flasks, incubated at 37°C, 200 rpm for 16 hours. These single clone cultures were used for predation assays and genome sequencing.

### Predation assay

To evaluate predation resistance and growth dynamics of different prey, predation assays were performed. Different *E. coli* prey strains were assessed: *E. coli* K-12 MG1655 population lineages evolved under high (H1-8), low (L1-8) and no (N1-8) predatory pressure at the end 15^th^ evolution cycle (after recovery phase) were compared with ancestor *E. coli* K-12 MG1655. At first, we exposed prey lineages that evolved under a particular predatory pressure to that pressure. We then focussed on exposing evolved lines to the high predatory pressure condition.

To investigate the temporal aspect of evolution, we performed predation assays of prey lineages evolved under high predatory pressure (H1-H8) at the end of the 2^nd^ and 5^th^ evolution cycle and compared them to the 15^th^ cycle, when exposed to high predatory pressure. To test for cross resistance, we assessed the performance of prey that evolved under no or low predator by exposing them to high predatory pressure. The predation assays at the population level featured 5 replicates per prey-predator combination and were repeated on two separate days.

To assess effects of specific prey genes on predation resistance, predation assay was conducted with markerless single gene deletion mutants (Δ*envZ,* Δ*ompF,* Δ*truA,* Δ*waaB,* and Δ*waaF* based on *E. coli* BW25113 (reference strain from KEIO library^2^). These strains were exposed to high and no predatory pressure conditions.

Prior to the predation assay, respective evolved prey lineages or prey strains (evolved lineages, population- or single clone-based, single gene deletion, and KEIO reference strain) were grown from cryo stocks by inoculation in 50 ml YT broth and incubation at 37°C and 200 rpm for 16 h. Additionally, an overnight culture of ancestor prey was prepared from a single colony, at same growth conditions. The eight independently evolved prey lineages under each predatory pressure were assessed in parallel alongside ancestor prey. 10-ml prey overnight cultures were centrifuged at 5311 x *g* for 3 min and subsequently washed with 10 ml of Ca-HEPES buffer. Then prey cells were resuspended in 6 ml of Ca-HEPES buffer and adjusted to OD_600nm =_ 0.03. A 24-hr predator lysate was filtered through a 0.8-μm syringe filter (Whatman® Puradisc FP 30, Cellulose Acetate, sterile, WHA10462240). For the predation assays, the same cultivation media was used as for the experimental evolution predation phase, comprising 15 ml of YT broth, 5 ml of 25 mM HEPES and 2 mM CaCl_2 o_verall. We set up the predation assay in 96-well plates, in a total volume of 150 μl. Initially 119 μl of cultivation media was preset, and different volumes of the filtered predator lysate were added to the assay: 0 μl (no predator exposure), 4 μl (low predator exposure) or 16 μl (high predator exposure), depending on the evolved prey lineages and predatory pressure tested. Then we added 15 μl of the respective prey solution (OD_600nm =_ 0.03 in Ca-HEPES). Then we adjusted the final volume in reaction wells to 150 μl by adding Ca-HEPES (as proxy for predator lysate and prey resuspension), resulting in a final concentration of prey of OD_600nm =_ 0.003. We used the same ratio of prey and predator as in the predation phase of the prey evolution experiment. The plate was incubated in a plate reader (Tecan Infinite 200 Pro) at 29°C for 24 h. OD_600nm o_f the reaction wells was measured with 10 flashes every 5 minutes after shaking at 6-mm amplitude for 120 secs.

We used OD_600nm a_s proxy for the biomass of prey cells, as the predator cells only contribute to a very minor fraction to OD_600nm b_ased on the small size. The kinetic measurement of OD_600nm o_ver 24 h, was analysed using an interactive platform for analysis of microbial growth data (https://statu.shinyapps.io/ExploreMicrobialGrowth/, version 2016.01.12) designed by Adin Ross-Gillespie at University of Zurich. We used Spline function to fit the kinetic growth data to extract the integral (area under the curve) and maximal growth increase rate. For the predation assays in cultivation media, we derived relative growth integral and growth rate of evolved prey lineages in comparison to the ancestor prey. For this we divided the growth integral or growth rate of evolved lineages under all predator conditions by that of ancestor when exposed to no predator, that is baseline. Values > 1 or < 1, indicate that evolved lineages perform better or worse than baseline.

### Cell shape analysis by phase-contrast microscopy

*E. coli* K-12 lineages evolved under low, no, and high predatory pressure at the end of experimental evolution (after the 15^th^ evolution cycle), *E. coli* K-12 MG1655 (ancestor), as well as *E. coli* K-12 BW25113, Δ*envZ* and Δ*truA* were regrown from cryo stocks in YT broth at 37°C and 200 rpm shaking overnight. From each culture 10 µl was applied onto a 1% agarose in Ca/HEPES buffer pad on a clean microscope slide for phase-contrast microscopy. Imaging was performed using a Leica DMi8 inverted microscope equipped with a 100x objective (HC PL APO 1.40 oil PH3), a Leica monochrome fluorescence DFC9000 gTC Camera System, and Leica Application Suite X Software Version 3.7.6.25997 (Leica microsystems). Exposure time was 100 ms with illumination intensity of 140. For each evolved lineage and ancestor strain *E. coli* K-12 MG1655 a minimum of 15 images were collected. For shape analysis of the evolved cells, software Fjij/ImageJ (Version 2.90/1.53t, Java 1.8.0322, 64-bit)^6,7^ was used with the Plugin MicrobeJ (Version 5.13o (16) – beta)^8^. Area and length restrictions were set to 0.5-3 p^2^, width to 0.5-2 p^2^. After initial shape detection, masks were manually adjusted to observed cell size and a minimum of 300 cells per lineage/strain were analysed.

### Statistical analyses

We used general linear models for statistical analysis in R 4.2.3^9,10^. To assess whether relative growth integral and relative growth rate vary in response to predatory pressure we used analysis of variance (ANOVA) models (Fig. 1e-g, Fig. 2 c,d & Fig. 3d-f). We further used two-sided one-sample *t*-tests to test for significant differences in relative growth integral or growth rate between evolved prey lineages and that of ancestor when exposed to either no, low or high predatory pressure (i.e. being different from mean relative growth integral or growth rate of ancestor, Fig. 1e-g & Fig. 2c-d & Fig. 3d-f & Fig. 5b,d & Extended Data Fig. 2d-f, j-l). Reported p-values for *t-*tests were adjusted for multiple testing using the Holm method. We used a logistic regression model to test for a negative association (tradeoff) between mean relative growth integral and mean relative growth rate (Fig. 3g). A detailed statistical output of the data of this article is available in an Additional Online Material table.

### Genome sequencing and analysis of evolved prey population lineages

We sequenced the genome of the 24 prey lineages (i.e. population) that evolved under no, low and high predatory pressure condition (8 each) after the 15^th^ evolution cycle, at the population level and compared it with the ancestor prey genome. Overnight cultures were grown in YT broth (200 rpm at 37°C) from cryo stocks of evolved prey lineages from the 15^th^ recovery phase at the end of experimental evolution and separately from a single colony of the ancestor prey. From these cultures we extracted genomic DNA using GenElute^TM^ Bacterial Genomic DNA Kit (Sigma, NA2110). The concentration of genomic DNA varied between 83.77 to 205. 407 ng/μl, which was sufficient for standard library preparation. The samples were then transferred to the Functional Genomic Center Zurich (FGCZ, Switzerland) for sequencing and base variant calling of the sequencing data of evolved lineages against the ancestor. For library preparation the TruSeq DNA Nano Sample Prep Kit v2 (Illumina, Inc, California, USA) was used in the succeeding steps. DNA samples (100ng) were sonicated with the Covaris using settings specific to the fragment size of 350 bp. The fragmented DNA samples were size-selected using AMpure beads, end-repaired and adenylated. TruSeq adapters containing Unique Dual Indices (UDI) for multiplexing were ligated to the size-selected DNA samples. Fragments containing TruSeq adapters on both ends were selectively enriched by PCR. The quality and quantity of the enriched libraries were validated using TapeStation (Agilent, Waldbronn, Germany). An average fragment size of approximately 500 bp was achieved. The libraries were normalized to 10 nM in Tris-Cl 10 mM, pH 8.5 with 0.1% Tween 20. For sequencing and cluster generation the Novaseq 6000 (Illumina, Inc, California, USA) was used according to standard protocol. Sequencing were paired end at 2 X150 bp. Technical quality of Illumina paired-end (PE) reads was evaluated using FastQC (v0.11.9) and FastQScreen (v0.14.1). Raw reads were pre-processed using fastp (v0.20.0) with the following parameter: hard trimming of 10 nucleotide (nt) and 1 nt at 5’ and 3 ends, respectively; quality trimming of both ends using a sliding window of 4 nt and mean quality of 20; trimmed reads shorter than 18 nt, and trimmed reads with mean read quality lower than 20 were filtered out. Pre-processed PE reads were aligned to the reference genome (*E. coli* K-12 MG1655 refseq GCF_000005845.2) using bwa (v0.7.17) with the “mem” algorithm. PCR duplicates were marked using Picard (v2.22.8, MarkDuplicates). Mapping results were quality controlled using SAMStat (v1.5.1), Qualimap (v2.2.1), and Picard (v2.22.8, CollectGcBiasMetrics). Multi-sample variant calling was performed using freebayes (V1.3.2) with frequency cutoffs of 5% and 1%, respectively. Only unique reads in primary alignments with alignment quality 30 and above, and bases with quality 20 and above, were counted for variant analysis. Sites with multiple alternative alleles were split into multiple rows using bcftools. Variant annotation was performed using snpEff (v4.3) against reference gene models (NCBI, reference genome: ASM584v2, refseq: GCF_000005845.2). Called variants were filtered against the ancestor strain using a customized Perl script. Markers were retained, where all counted reads contain only the reference alleles in the ancestor strain. Per sample variant frequencies were then calculated for these markers using the customized Perl script. The resulting excel file from population variant calling was used for further analysis and representation by hierarchically clustered heatmaps on absolute mutation frequencies and their differences compared to the ancestor strain. Heatmaps were generated in R with Pheatmap^11^. We focused on gene mutations with a frequency more than 5%. If a particular gene had multiple mutation effects, we selected the mutation with the highest frequency. Mutation counts were added to a gene only if the mutation was located within the coding region of a specific gene (direct effects). This excludes downstream effects on adjacent genes. Mutations lying in intergenic regions were ignored due to their ambiguous impact on surrounding genes. An exception was made for the mutation located in the intergenic region between *asnS* and *ompF*. Since it is located 143 bp upstream the *ompF* transcription start site various *ompF* regulators bind adjacent to this locus and is likely to influence *ompF* transcription. The mutations in *ompF* were analysed in detail and the OmpF amino acid sequence with changes in the different prey lineages was constructed in Excel (Microsoft Corporation, 2018) and compared with the ancestor strain. Uniquely mutated genes (without transposases and rRNA genes) of all genomes of prey lineages at the respective predatory pressure were plotted using R package VennDiagram^12^.

### Genome sequencing and variant calling for prey single clones

48 cultures were prepared from six single prey clones each of all eight prey lineages evolved under high predatory pressure (H1-H8) at the end of the 15^th^ evolution cycle. Overnight cultures of these single clones from lineages evolved under high-predatory pressure were inoculated from cryo stocks into 10 ml YT medium in 50 ml Falcon tubes, and incubated at 37°C, 200 rpm for 16 h. We harvested prey cells equivalent of OD_600nm =_ 10.0 from these cultures by centrifugation at 5000 *g* for 5 min, the cells were then washed (5000 g for 5 minutes) in 1 ml Phosphate buffered saline (Gibco) and resuspended into 0.5ml DNA/RNA shield buffer (Zymo Research). This mixture was sent to MicrobesNG (UK, https://microbesng.com, Birmingham, UK), for DNA extraction, sequencing and base variant calling against reference genome *E. coli* K-12 MG1655.

DNA was extracted by lysing 4-5 µl cell suspension with 120 μL of TE buffer containing lysozyme (MPBio, USA), metapolyzyme (Sigma Aldrich, USA) and RNase A (ITW Reagents, Spain), and incubated for 25 min at 37°C. Proteinase K (VWR Chemicals, Ohio, USA) (final concentration 0.1mg/mL) and SDS (Sigma-Aldrich, Missouri, USA) (final concentration 0.5% v/v) are added and incubated for 5 min at 65°C. Genomic DNA was purified using an equal volume of SPRI beads and resuspended in EB buffer (10mM Tris-HCl, pH 8.0). The DNA was quantified with the Quant-iT dsDNA HS (ThermoFisher Scientific) assay in an Eppendorf AF2200 plate reader (Eppendorf UK Ltd, United Kingdom) and diluted as appropriate. Genomic DNA libraries are prepared using the Nextera XT Library Prep Kit (Illumina, San Diego, USA) following the manufacturer’s protocol with the following modifications: input DNA is increased 2-fold, and PCR elongation time is increased to 45 sec. DNA quantification and library preparation are carried out on a Hamilton Microlab STAR automated liquid handling system (Hamilton Bonaduz AG, Switzerland). Libraries are sequenced on an lllumina NovaSeq 6000 (Illumina, San Diego, USA) using a 250 bp paired end protocol. For projects predating April, 2020, libraries were sequenced on an Illumina HiSeq 2500 using a 250 bp paired-end protocol (Illumina, San Diego, USA). Reads are adapter trimmed using Trimmomatic version 0.30 ^13^ with a sliding window quality cutoff of Q15. De novo assembly is performed on samples using SPAdes version 3.7 ^14^, and contigs are annotated using Prokka version 1.11 ^15^. Reads were aligned to the reference using BWA mem v0.7.17 ^16^ and are then processed using SAMtools v1.9 ^17^. Variants were called using VarScan v2.4.0 ^18^ with two thresholds, sensitive and specific, where the variant allele frequency is greater than 10% respectively. The effects of the variants are predicted and annotated using SnpEff v4.3 ^19^. The base variant calling was performed using the criteria of a sequence coverage of at least 3x and a frequency of at least 10% to generate a table reporting on the differences to the reference genome (NCBI, refseq: GCF_000005845.2). The resulting Microsoft Excel table obtained from Microbes NG was used to combine it with the population sequencing data based on the nucleotide position. Gene annotations were obtained from both population and clone sequencing data, with frequency more than 5%.

## Data availability

The raw data (OD_600nm m_easurements over time, microscopy images and raw data on width and length, R code used for analysis) used to generate the findings of this project have been deposited in the figshare repository (doi: XXX). The sequencing data for this study has been deposited in the European Nucleotide Archive (ENA) at EMBL-EBI under accession number PRJEB78919 (https://www.ebi.ac.uk/ena/browser/view/PRJEB78919).

## References (Main text)

1. Murray, C. J., et al. Global burden of bacterial antimicrobial resistance in 2019: a systematic analysis. The Lancet 399, 629–655 (2022).

2. Pirnay, J.-P., et al. Personalized bacteriophage therapy outcomes for 100 consecutive cases: a multicentre, multinational, retrospective observational study. Nat Microbiol 9, 1434–1453 (2024).

3. Sockett, R. E. & Lambert, C. Bdellovibrio as therapeutic agents: a predatory renaissance? Nat Rev Microbiol 2, 669–675 (2004).

4. Konwar, A. N., Hazarika, S. N., Bharadwaj, P. & Thakur, D. Emerging Non-Traditional Approaches to Combat Antibiotic Resistance. Curr Microbiol 79, 330 (2022).

5. Koskella, B., Hernandez, C. A. & Wheatley, R. M. Understanding the Impacts of Bacteriophage Viruses: From Laboratory Evolution to Natural Ecosystems. Annu. Rev. Virol. 9, 57–78 (2022).

6. Atterbury, R. J. & Tyson, J. Predatory bacteria as living antibiotics - where are we now? Microbiology (Reading) 167, (2021).

7. Lai, T. F., Ford, R. M. & Huwiler, S. G. Advances in cellular and molecular predatory biology of *Bdellovibrio bacteriovorus* six decades after discovery. Front. Microbiol. 14, 1168709 (2023).

8. Saralegui, C., Herencias, C., Halperin, A. V., De Dios-Caballero, J., Pérez-Viso, B., Salgado, S., Lanza, V. F., Cantón, R., Baquero, F., Prieto, M. A. & Del Campo, R. Strain-specific predation of *Bdellovibrio bacteriovorus* on *Pseudomonas aeruginosa* with a higher range for cystic fibrosis than for bacteremia isolates. Sci Rep 12, 10523 (2022).

9. Upatissa, S., Mun, W. & Mitchell, R. J. Pairing Colicins B and E5 with *Bdellovibrio bacteriovorus* To Eradicate Carbapenem- and Colistin-Resistant Strains of *Escherichia coli*. Microbiol Spectr 11, e00173–23 (2023).

10. Sun, Y., Ye, J., Hou, Y., Chen, H., Cao, J. & Zhou, T. Predation Efficacy of *Bdellovibrio bacteriovorus* on Multidrug-Resistant Clinical Pathogens and Their Corresponding Biofilms. Jpn J Infect Dis 70, 485–489 (2017).

11. Dashiff, A., Junka, R. A., Libera, M. & Kadouri, D. E. Predation of human pathogens by the predatory bacteria *Micavibrio aeruginosavorus* and *Bdellovibrio bacteriovorus*: Predation by *M. aeruginosavorus* and *B. bacteriovorus*. Journal of Applied Microbiology 110, 431–444 (2011).

12. Jurkevitch, E., Minz, D., Ramati, B. & Barel, G. Prey Range Characterization, Ribotyping, and Diversity of Soil and Rhizosphere *Bdellovibrio* spp. Isolated on Phytopathogenic Bacteria. Appl Environ Microbiol 66, 2365–2371 (2000).

13. Im, H., Choi, S. Y., Son, S. & Mitchell, R. J. Combined Application of Bacterial Predation and Violacein to Kill Polymicrobial Pathogenic Communities. Sci Rep 7, 14415 (2017).

14. Koval, S. F. & Hynes, S. H. Effect of paracrystalline protein surface layers on predation by *Bdellovibrio bacteriovorus*. J Bacteriol 173, 2244–2249 (1991).

15. Mun, W., Upatissa, S., Lim, S., Dwidar, M. & Mitchell, R. J. Outer Membrane Porin F in *E. coli* Is Critical for Effective Predation by *Bdellovibrio*. Microbiol Spectr 10, e03094–22 (2022).

16. Georjon, H. & Bernheim, A. The highly diverse antiphage defence systems of bacteria. Nat Rev Microbiol 21, 686–700 (2023).

17. Varon, M. Selection of predation-resistant bacteria in continuous culture. Nature 277, 386–388 (1979).

18. Gallet, R., Alizon, S., Comte, P.-A., Gutierrez, A., Depaulis, F., van Baalen, M., Michel, E. & Muller-Graf, C. D. M. Predation and Disturbance Interact to Shape Prey Species Diversity. 170, 143–154 (2007).

19. Gallet, R., Tully, T. & Evans, M. E. K. Ecological conditions affect evolutionary trajecory in a predator-prey system. Evolution 63, 641–651 (2009).

20. Caulton, S. G., Lambert, C., Tyson, J., Radford, P., Al-Bayati, A., Greenwood, S., Banks, E. J., Clark, C., Till, R., Pires, E., Sockett, R. E. & Lovering, A. L. *Bdellovibrio bacteriovorus* uses chimeric fibre proteins to recognize and invade a broad range of bacterial hosts. Nat Microbiol 9, 214–227 (2024).

21. Chanyi, R. M., Ward, C., Pechey, A. & Koval, S. F. To invade or not to invade: two approaches to a prokaryotic predatory life cycle. Can J Microbiol 59, 273–279 (2013).

22. Shemesh, Y. & Jurkevitch, E. Plastic phenotypic resistance to predation by *Bdellovibrio* and like organisms in bacterial prey. Environmental Microbiology 6, 12–18 (2004).

23. Aharon, E., Mookherjee, A., Pérez-Montaño, F., Mateus Da Silva, G., Sathyamoorthy, R., Burdman, S. & Jurkevitch, E. Secretion systems play a critical role in resistance to predation by *Bdellovibrio bacteriovorus*. Research in Microbiology 172, 103878 (2021).

24. Hoshiko, Y., Nishiyama, Y., Moriya, T., Kadokami, K., López-Jácome, L. E., Hirano, R., García-Contreras, R. & Maeda, T. Quinolone Signals Related to Pseudomonas Quinolone Signal-Quorum Sensing Inhibits the Predatory Activity of *Bdellovibrio bacteriovorus*. Front. Microbiol. 12, 722579 (2021).

25. Dwidar, M., Jang, H., Sangwan, N., Mun, W., Im, H., Yoon, S., Choi, S., Nam, D. & Mitchell, R. J. Diffusible Signaling Factor, a Quorum-Sensing Molecule, Interferes with and Is Toxic Towards *Bdellovibrio bacteriovorus* 109J. Microb Ecol 81, 347–356 (2021).

26. Dwidar, M., Nam, D. & Mitchell, R. J. Indole negatively impacts predation by *B dellovibrio bacteriovorus* and its release from the bdelloplast. Environmental Microbiology 17, 1009–1022 (2015).

27. Mun, W., Kwon, H., Im, H., Choi, S. Y., Monnappa, A. K. & Mitchell, R. J. Cyanide Production by *Chromobacterium piscinae* Shields It from *Bdellovibrio bacteriovorus* HD100 Predation. mBio 8, e01370–17 (2017).

28. Andersson, D. I. & Hughes, D. Antibiotic resistance and its cost: is it possible to reverse resistance? Nat Rev Microbiol 8, 260–271 (2010).

29. Andersson, D. I. The biological cost of mutational antibiotic resistance: any practical conclusions? Current Opinion in Microbiology 9, 461–465 (2006).

30. Cai, S. J. & Inouye, M. EnvZ-OmpR Interaction and Osmoregulation in *Escherichia coli*. Journal of Biological Chemistry 277, 24155–24161 (2002).

31. Davis, D. R. & Poulter, C. D. Proton-nitrogen-15 NMR studies of *Escherichia coli* tRNAPhe from HisT mutants: a structural role for pseudouridine. Biochemistry 30, 4223–4231 (1991).

32. Tsui, H. C., Arps, P. J., Connolly, D. M. & Winkler, M. E. Absence of hisT-mediated tRNA pseudouridylation results in a uracil requirement that interferes with *Escherichia coli* K-12 cell division. J Bacteriol 173, 7395–7400 (1991).

33. Gronow, S., Brabetz, W. & Brade, H. Comparative functional characterization *in vitro* of heptosyltransferase I (WaaC) and II (WaaF) from *Escherichia coli*. European Journal of Biochemistry 267, 6602–6611 (2000).

34. Aminov, R. I. A Brief History of the Antibiotic Era: Lessons Learned and Challenges for the Future. Front. Microbio. 1, (2010).

35. Davies, J. & Davies, D. Origins and Evolution of Antibiotic Resistance. Microbiol Mol Biol Rev 74, 417–433 (2010).

36. Fang, Q., Feng, Y., McNally, A. & Zong, Z. Characterization of phage resistance and phages capable of intestinal decolonization of carbapenem-resistant *Klebsiella pneumoniae* in mice. Commun Biol 5, 48 (2022).

37. Oechslin, F. Resistance Development to Bacteriophages Occurring during Bacteriophage Therapy. Viruses 10, 351 (2018).

38. Maffei, E., Shaidullina, A., Burkolter, M., Heyer, Y., Estermann, F., Druelle, V., Sauer, P., Willi, L., Michaelis, S., Hilbi, H., Thaler, D. S. & Harms, A. Systematic exploration of *Escherichia coli* phage–host interactions with the BASEL phage collection. PLoS Biol 19, e3001424 (2021).

39. Nobrega, F. L., Vlot, M., De Jonge, P. A., Dreesens, L. L., Beaumont, H. J. E., Lavigne, R., Dutilh, B. E. & Brouns, S. J. J. Targeting mechanisms of tailed bacteriophages. Nat Rev Microbiol 16, 760–773 (2018).

40. Jochumsen, N., Marvig, R. L., Damkiær, S., Jensen, R. L., Paulander, W., Molin, S., Jelsbak, L. & Folkesson, A. The evolution of antimicrobial peptide resistance in *Pseudomonas aeruginosa* is shaped by strong epistatic interactions. Nat Commun 7, 13002 (2016).

41. Kenney, L. J. & Anand, G. S. EnvZ/OmpR Two-Component Signaling: An Archetype System That Can Function Noncanonically. EcoSal Plus 9, 10.1128/ecosalplus.ESP-0001-2019 (2020).

42. Thanassi, D. G., Suh, G. S. & Nikaido, H. Role of outer membrane barrier in efflux-mediated tetracycline resistance of *Escherichia coli*. J Bacteriol 177, 998–1007 (1995).

43. Choi, U. & Lee, C.-R. Distinct Roles of Outer Membrane Porins in Antibiotic Resistance and Membrane Integrity in *Escherichia coli*. Front. Microbiol. 10, 953 (2019).

44. Delcour, A. H. Outer membrane permeability and antibiotic resistance. Biochimica et Biophysica Acta (BBA) - Proteins and Proteomics 1794, 808–816 (2009).

45. Tyson, J., Radford, P., Lambert, C., Till, R., Huwiler, S. G., Lovering, A. L. & Elizabeth Sockett, R. Prey killing without invasion by *Bdellovibrio bacteriovorus* defective for a MIDAS-family adhesin. Nat Commun 15, 3078 (2024).

46. Varon, M. & Shilo, M. Attachment of *Bdellovibrio bacteriovorus* to cell wall mutants of *Salmonella* spp. and *Escherichia coli*. J Bacteriol 97, 977–979 (1969).

47. Schelling, M. & Conti, S. Host receptor sites involved in the attachment of *Bdellovibrio bacteriovorus* and *Bdellovibrio stolpii*. FEMS Microbiology Letters 36, 319–323 (1986).

48. Duncan, M. C., Forbes, J. C., Nguyen, Y., Shull, L. M., Gillette, R. K., Lazinski, D. W., Ali, A., Shanks, R. M. Q., Kadouri, D. E. & Camilli, A. *Vibrio cholerae* motility exerts drag force to impede attack by the bacterial predator *Bdellovibrio bacteriovorus*. Nat Commun 9, 4757 (2018).

49. Wale, N., Sim, D. G., Jones, M. J., Salathe, R., Day, T. & Read, A. F. Resource limitation prevents the emergence of drug resistance by intensifying within-host competition. Proc. Natl. Acad. Sci. U.S.A. 114, 13774–13779 (2017).

## References (Methods)

1. Lambert, C. & Sockett, R. E. Laboratory maintenance of Bdellovibrio. Curr Protoc Microbiol Chapter 7, Unit 7B.2 (2008).

2. Baba, T., Ara, T., Hasegawa, M., Takai, Y., Okumura, Y., Baba, M., Datsenko, K. A., Tomita, M., Wanner, B. L. & Mori, H. Construction of *Escherichia coli* K-12 in-frame, single-gene knockout mutants: the Keio collection. Molecular Systems Biology 2, 2006.0008 (2006).

3. Datsenko, K. A. & Wanner, B. L. One-step inactivation of chromosomal genes in *Escherichia coli* K-12 using PCR products. Proc. Natl. Acad. Sci. U.S.A. 97, 6640–6645 (2000).

4. Barrick Lab. Barrick Lab :: ProtocolsElectrocompetentCells. at <https://barricklab.org/twiki/bin/view/Lab/ProtocolsElectrocompetentCells>

5. Barrick Lab. Barrick Lab :: ProcedureFLPFRTRecombination. at <https://barricklab.org/twiki/bin/view/Lab/ProcedureFLPFRTRecombination>

6. Schindelin, J., Arganda-Carreras, I., Frise, E., Kaynig, V., Longair, M., Pietzsch, T., Preibisch, S., Rueden, C., Saalfeld, S., Schmid, B., Tinevez, J.-Y., White, D. J., Hartenstein, V., Eliceiri, K., Tomancak, P. & Cardona, A. Fiji: an open-source platform for biological-image analysis. Nat Methods 9, 676–682 (2012).

7. Rueden, C. T., Schindelin, J., Hiner, M. C., DeZonia, B. E., Walter, A. E., Arena, E. T. & Eliceiri, K. W. ImageJ2: ImageJ for the next generation of scientific image data. BMC Bioinformatics 18, 529 (2017).

8. Ducret, A., Quardokus, E. M. & Brun, Y. V. MicrobeJ, a tool for high throughput bacterial cell detection and quantitative analysis. Nat Microbiol 1, 16077 (2016).

9. R Core Team. R: A language and environment for statistical computing. (Vienna, Austria, 2021). at <https://www.R-project.org/>

10. Wickham, H., Averick, M., Bryan, J., Chang, W., McGowan, L., François, R., Grolemund, G., Hayes, A., Henry, L., Hester, J., Kuhn, M., Pedersen, T., Miller, E., Bache, S., Müller, K., Ooms, J., Robinson, D., Seidel, D., Spinu, V., Takahashi, K., Vaughan, D., Wilke, C., Woo, K. & Yutani, H. Welcome to the Tidyverse. JOSS 4, 1686 (2019).

11. Kolde, R. Pheatmap: pretty heatmaps. (R Package Version 1.0.10., 2019). at <https://CRAN.R-project.org/package=pheatmap>

12. Chen, H. VennDiagram: Generate High-Resolution Venn and Euler Plots. (R package version 1.7.3, 2022). at <https://CRAN.R-project.org/package=VennDiagram>

13. Bolger, A. M., Lohse, M. & Usadel, B. Trimmomatic: a flexible trimmer for Illumina sequence data. Bioinformatics 30, 2114–2120 (2014).

14. Bankevich, A., Nurk, S., Antipov, D., Gurevich, A. A., Dvorkin, M., Kulikov, A. S., Lesin, V. M., Nikolenko, S. I., Pham, S., Prjibelski, A. D., Pyshkin, A. V., Sirotkin, A. V., Vyahhi, N., Tesler, G., Alekseyev, M. A. & Pevzner, P. A. SPAdes: A New Genome Assembly Algorithm and Its Applications to Single-Cell Sequencing. Journal of Computational Biology 19, 455–477 (2012).

15. Seemann, T. Prokka: rapid prokaryotic genome annotation. Bioinformatics 30, 2068–2069 (2014).

16. Li, H. Aligning sequence reads, clone sequences and assembly contigs with BWA-MEM. Preprint at http://arxiv.org/abs/1303.3997 (2013)

17. Danecek, P., Bonfield, J. K., Liddle, J., Marshall, J., Ohan, V., Pollard, M. O., Whitwham, A., Keane, T., McCarthy, S. A., Davies, R. M. & Li, H. Twelve years of SAMtools and BCFtools. GigaScience 10, giab008 (2021).

18. Koboldt, D. C., Chen, K., Wylie, T., Larson, D. E., McLellan, M. D., Mardis, E. R., Weinstock, G. M., Wilson, R. K. & Ding, L. VarScan: variant detection in massively parallel sequencing of individual and pooled samples. Bioinformatics 25, 2283–2285 (2009).

19. Cingolani, P., Platts, A., Wang, L. L., Coon, M., Nguyen, T., Wang, L., Land, S. J., Lu, X. & Ruden, D. M. A program for annotating and predicting the effects of single nucleotide polymorphisms, SnpEff: SNPs in the genome of *Drosophila melanogaster* strain w ^1118^ ; iso-2; iso-3. Fly 6, 80–92 (2012).

