## Extended Data and Additional Online Material for "Outer membrane changes enable evolutionary escape from bacterial predation"

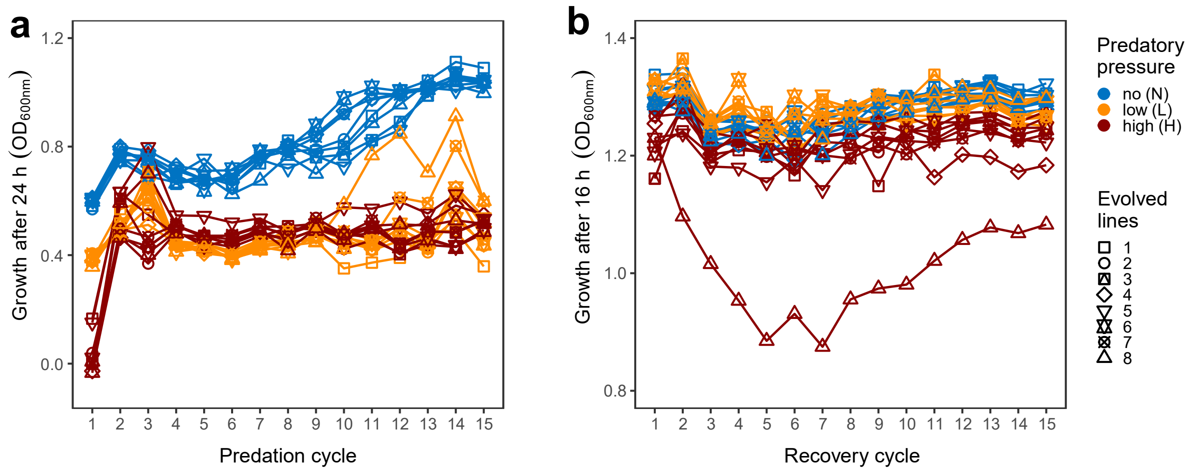


**Extended Data Fig. 1 | Growth of independently evolving prey lineages during experimental evolution.** A prey evolution experiment was conducted over 15 cycles, composed of an alternating predation and recovery phase, transferring only the evolved prey to the next evolution cycle (see Fig. **1a** for a detailed scheme). Evolved prey where then exposed to fresh predator in the subsequent predation phase. After separation of prey from predator, growth of prey in the absence of predator is defined as recovery phase. Growth was measured by $OD_{600nm}$ of eight independently evolved *E. coli* K-12 prey lineages (1-8, different shapes) when subjected to no (N, blue), low (L, orange) and high (H, red) predatory pressure. **a,** Growth trajectories over 15 evolution cycles measured at the end of each 24-h predation phase of *E. coli* K-12 prey lineages under different predatory pressure. **b,** Growth trajectories over the 15 evolution cycles measured at the end of each 16-h recovery phase.

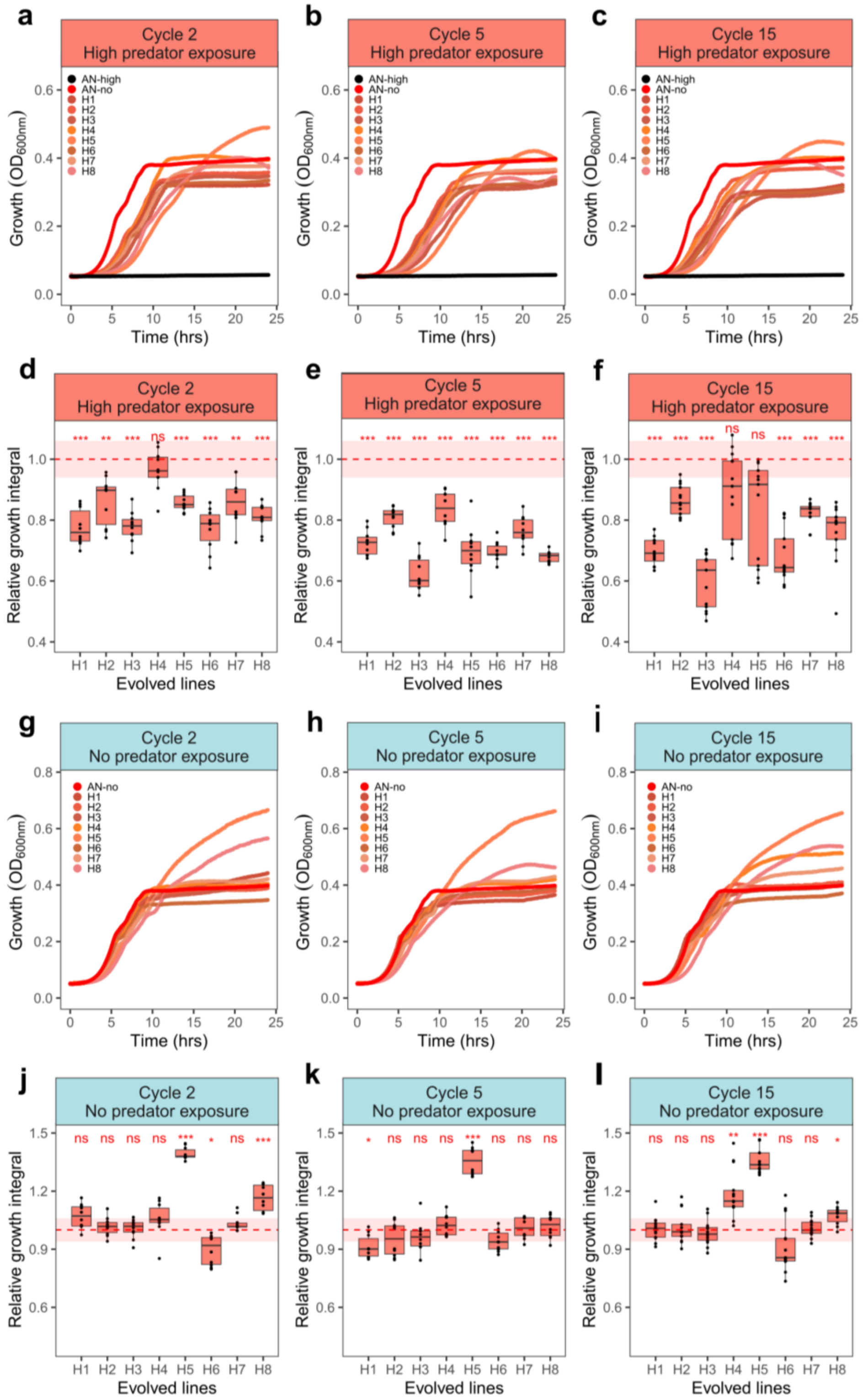


**Extended Data Fig. 2 │ Phenotypic predation resistance occurs early and remains consistent over the course of experimental prey evolution.** Growth of prey lineages evolved under high predatory pressure (H1-H8) was measured at the end of 2nd, 5th and 15th evolution cycle to determine temporal progression. As a reference the ancestor was exposed to no (AN-no, intense red) and high predatory pressure (AN-high, black). **a-c & g-i,** Growth kinetics measured as $OD_{600nm}$ when exposed to high **(a-c)** and no **(g-i)** predatory pressure. Each point of the growth curve represents the average $OD_{600nm}$ at that timepoint across replicates. **d-f** & **j-l,** Relative growth integral defined as growth integral (area under the curve) divided by the growth integral of ancestor under no predatory pressure. Red dotted line and shadow indicate the mean and standard deviation range of ancestor under no predation (AN-no). Boxplots represent the median with $25^{th}$ and $75^{th}$ percentiles, and whiskers show the 1.5 interquartile range. The p-value significance levels (two-sided one sample t-tests, $p>0.05$ = ns, $p<0.05$ = *, $p<0.01$ = **, $p<0.001$= ***) in comparison to AN-no is shown by red asterisks. Experiments were repeated independently at least twice, H1-8 $2^{nd}$ cycle (n= 10 replicates), H1-8 $5^{th}$ cycle (n= 10 replicates), H1-8 $15^{th}$ cycle (n= 13 replicates), AN-no (n= 72 replicates) and AN-high (n= 62 replicates). Plot **c** & **f** in this figure are identical to Fig.1 **d** & **g**, plot **i** is identical to Fig. **3c**. Precise p-values can be found in the extended online material (statistics table).

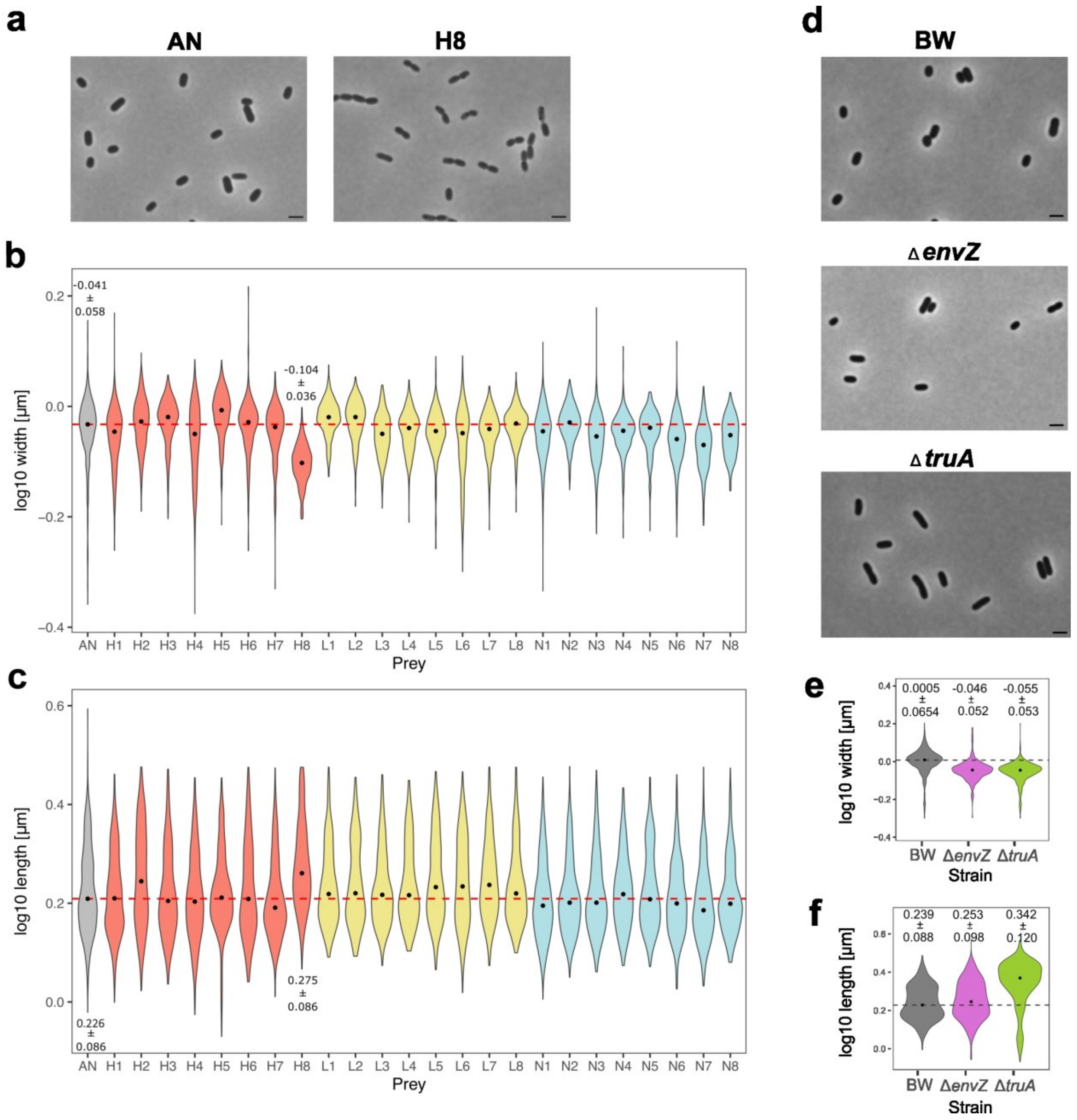


**Extended Data Fig. 3 │ Phase contrast microscopy revealed no differences in cell length and width between the ancestor and evolved prey population lineages, except for H8. Cell morphology changes in H8 are likely due to changes in *truA* (longer) and potentially *envZ* (slimmer). a & d,** Phase contrast microscopy images of ancestor *E. coli* K-12 (AN) and evolved *E. coli* K-12 prey population lineages evolved under high predatory pressure Nr. 8 (H8, **a**), as well as *E. coli* BW25113 (BW, KEIO reference strain) and clean-deletion mutants Δ*envZ* and Δ*truA* thereof (**d**). The scale bar is 2 µm. **b-c & e-f**, Violin plots show the quantitative analysis of microscopy images at single cell level for width (**b**) and length (**c**) of AN and eight independently evolved *E. coli* K-12 prey population lineages that evolved under high (H1-8), low (L1-8) and no (N1-8) predatory pressure for 15 evolution cycles. Same data for width (**e**) and length (**f**) for BW, Δ*envZ* and Δ*truA*. Numbers in the panels **b-c & e-f** indicate the mean and standard deviation for a particular population, black dot represent median of each population and dotted line represent the median of AN or BW. The following numbers of cells per lineage/ancestor (AN) were analysed: AN: 1516, H1: 735, H2: 684, H3: 596, H4: 1350, H5: 555, H6:483 , H7: 398, H8: 434, L1: 497, L2: 422, L3: 446 ,L4: 306, L5: 514, L6: 489, L7: 368, L8: 440, N1: 1565, N2: 877, N3: 467, N4: 399, N5: 588, N6: 437, N7: 582, N8: 397, BW: 869, Δ*envZ*: 872, Δ*truA*: 862. Mean, median and standard deviation of single cells for **b-c & e-f** is listed in additional online material

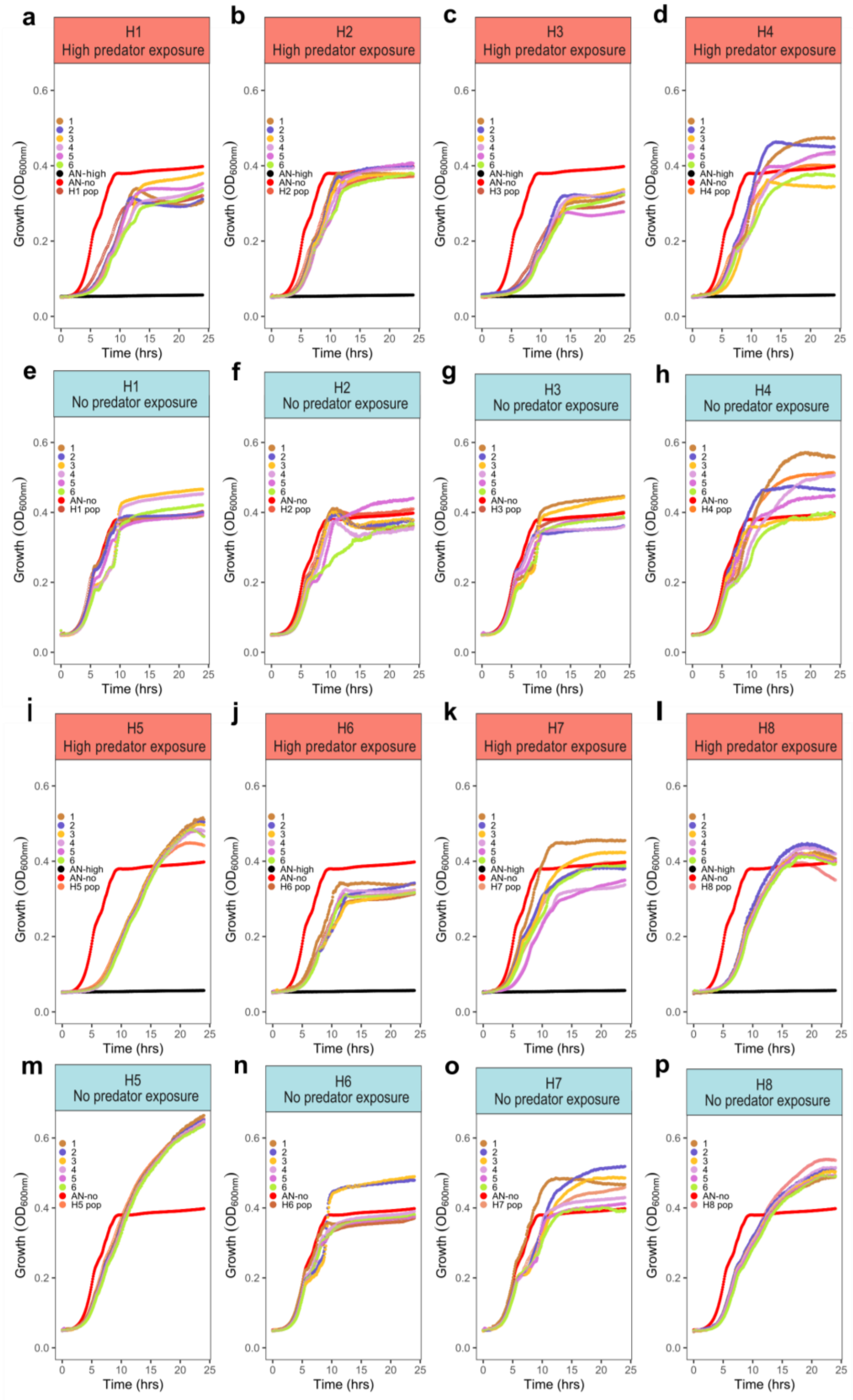


**Extended Data Fig. 4 │ Growth kinetics of single clones in comparison to populations that evolved under high predatory pressure and the ancestor.** Growth kinetics measured as $OD_{600nm}$ of single clones of high predatory pressure evolved lineages when exposed to **a-d, i-l**, high predatory pressure: H1 (**a**), H2 (**b**), H3 (**c**), H4 (**d**), H5 (**i**), H6 (**j**), H7 (**k**) and H8 (**l**), and to **e-h, m-p**, no predatory pressure: H1 (**e**), H2 (**f**), H3 (**g**), H4 (**h**), H5 (**m**), H6 (**n**), H7 (**o**)

and H8 (**p**), compared with ancestor (AN-no, red and AN-high, black). Each point of the growth curve represents the average $OD_{600nm}$ at that timepoint across replicates. Labels 1-6 denote single clones, H1-H8 denote populations that evolved under high predation pressure. H1-8 single clones (n= 3), H1-8 populations (n= 13), AN-no (n= 72) and AN-high (n= 62).

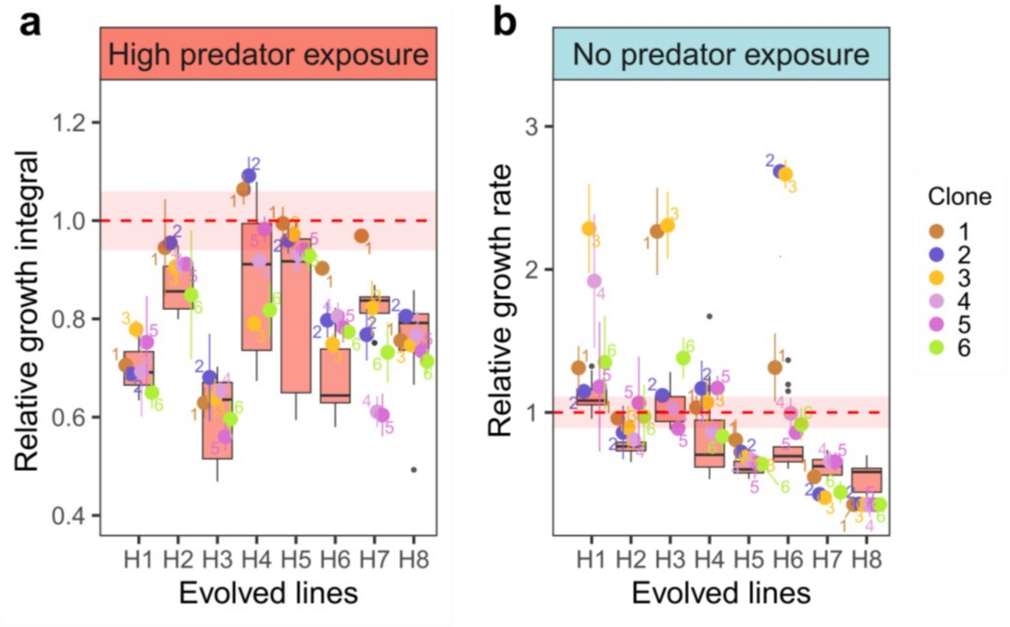


**Extended Data Fig. 5 │ Variation in resistance between evolved prey single clones and populations. a**, Relative growth integral, defined as growth integral (area under the curve) of the single clones or populations divided by the growth integral of ancestor under no predatory pressure and **b**, relative growth rate, defined as maximum growth rate of the single clones or populations divided by the maximum growth rate of the ancestor under no predatory pressure of single clones (Nr. 1-6 in different colours) of H1-H8 from 15$^{th}$ evolution cycle, under high and no predatory pressure respectively. Coloured circles represent the mean and coloured error bars represent the standard deviation for each clone. Boxplots represent the median with 25$^{th}$ and 75$^{th}$ percentiles for the evolved population, and whiskers show the 1.5 interquartile range. Black dots represent the outliers of H1-8 evolved population. Red dotted line and shadow indicate the mean and standard deviation range of ancestor under no predatory pressure (AN-no). H1-8 single clones (n= 3 replicates), H1-8 population (n= 13 replicates), AN-no (n= 72 replicates).

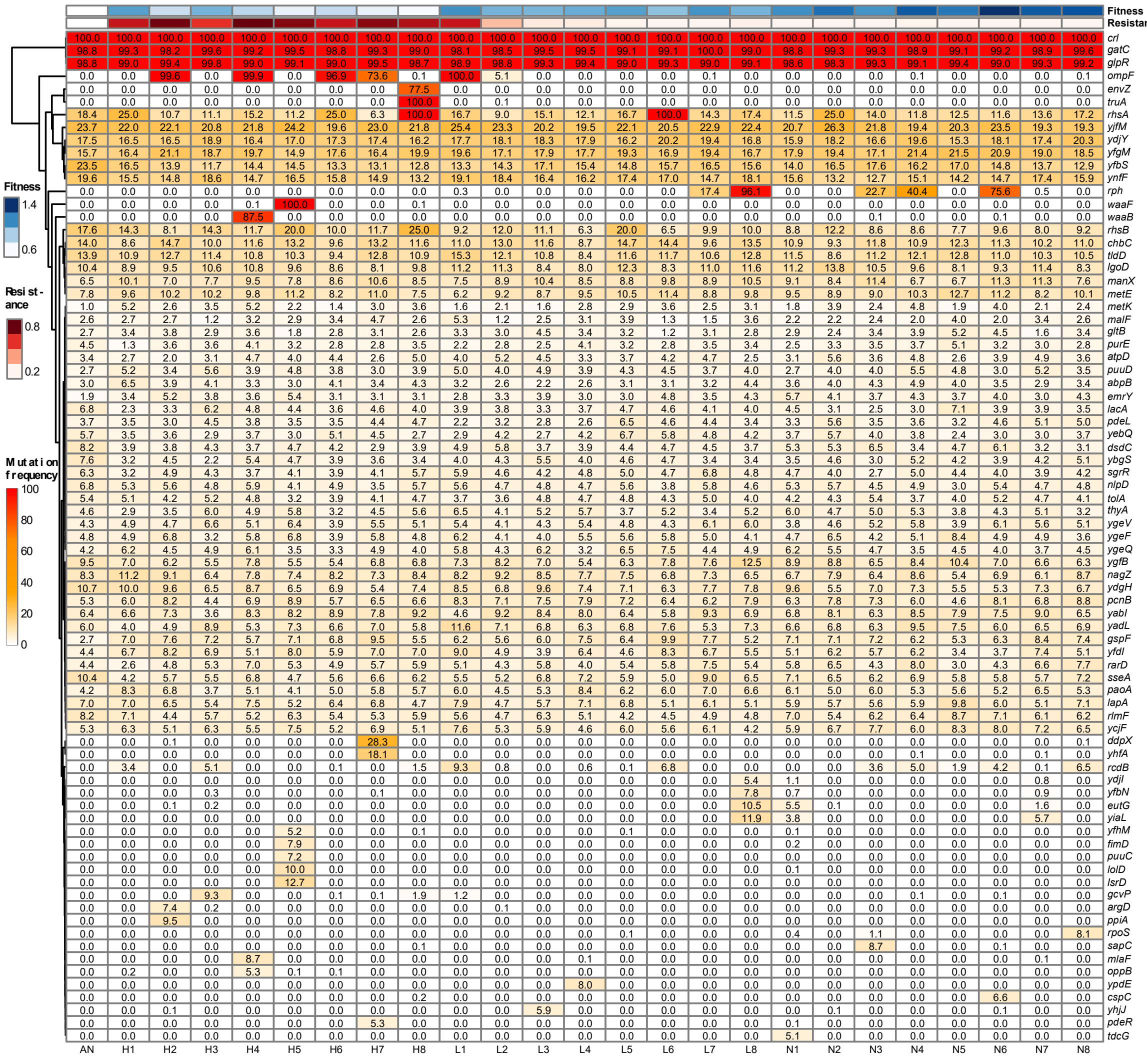


**Extended Data Fig. 6 │ Diverse genome-wide mutation patterns within genes of all evolved *E. coli* K-12 prey population lineages.** Hierarchically clustered heat map off all gene mutations with at least 5% frequency compared to *E. coli* K-12 MG1655 within a prey population lineage (excluding rRNA and transposase genes, in Extended Data Fig. S7) that evolved under high (H1-8), low (L1-8), and no (N1-8) predatory pressure at the 15$^{th}$ evolution cycle, as well as the ancestor (AN). Population fitness (blue heatmap, average relative growth rate of the lineages under no predatory pressure) and resistance to predator (red heatmap, average relative growth integral of the lineages under high predatory pressure) is indicated on top (see Fig. 3).

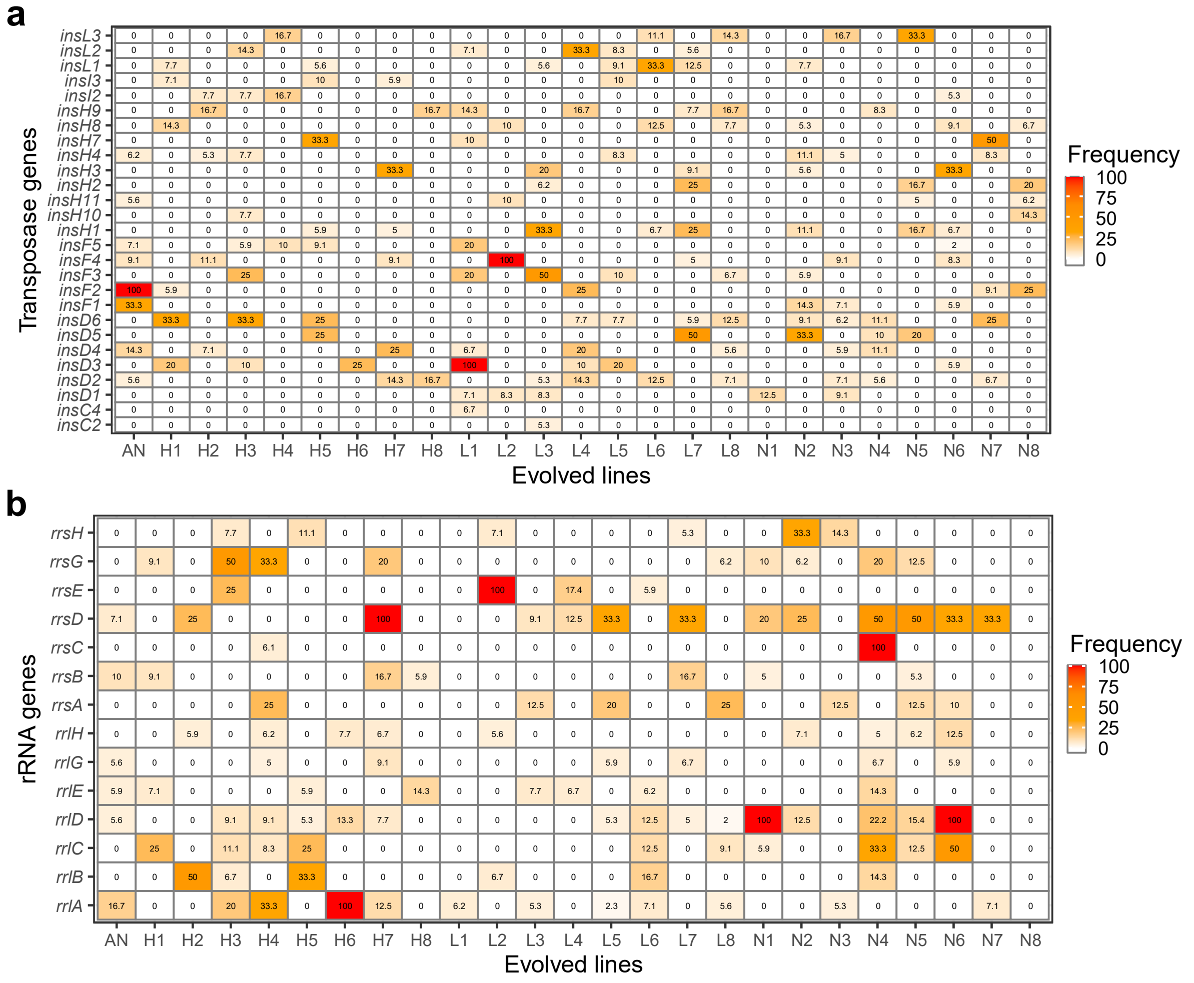


**Extended Data Fig. 7 │ Mutation frequency of the transposases and rRNA genes.** Heatmap with **a,** transposase and **b,** rRNA genes with at least 5% mutation frequency within a prey population that evolved under high (H1-8), low (L1-8), and no (N1-8) predatory pressure, at the 15$^{th}$ evolution cycle as well as ancestor (AN), mapped/compared against *E. coli* K-12 MG1655.

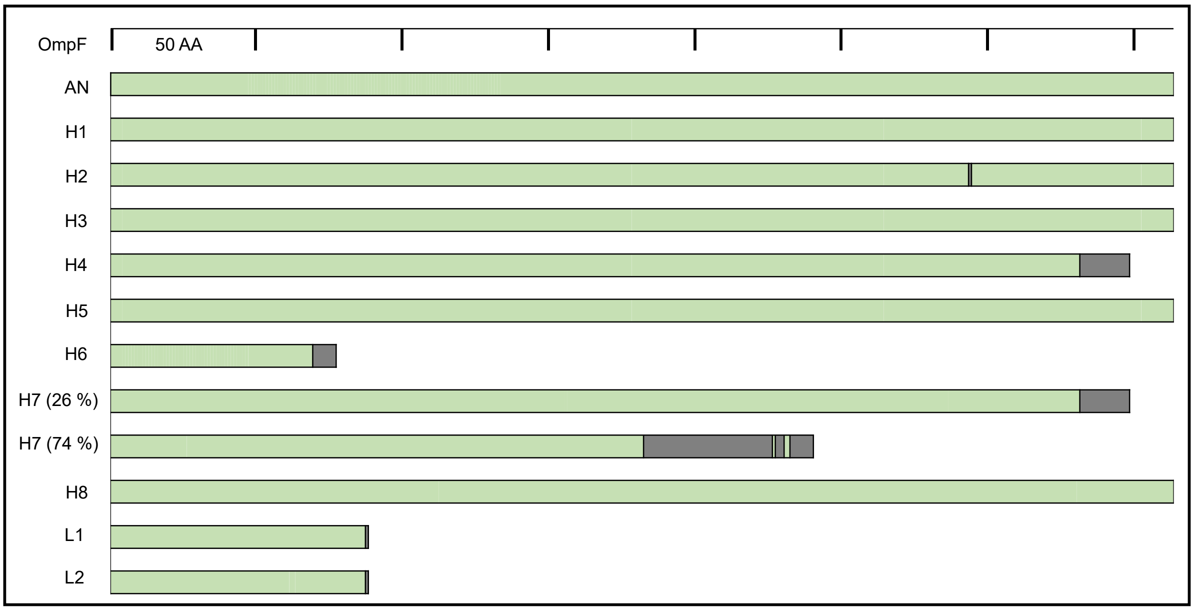


**Extended Data Fig. 8 │ Critical mutations occurring within the *ompF* gene resulting in the premature stop of OmpF in multiple evolved lineages.** Comparison of *ompF* gene of the *E. coli* K-12 MG1655 reference genome to the ancestor (AN), and different *E. coli* prey lineages evolved under high (H1-H8) and low predatory pressure (L1, L2). In lineage H3 no mutation within the coding sequence of *ompF*, but a point mutation in the intergenic region *ompF*-*asnS* upstream of *ompF* was detected (original C mutated to T at position 987125 at 100% of the population). Within lineage H7 two different types of mutations occurred, at different % within the population. Green bars represent identical translated amino acid (AA) sequences, grey bar represent different mutations at a specific locus, in many cases leading to a premature termination of gene translation (details in Extended Data Table 1).

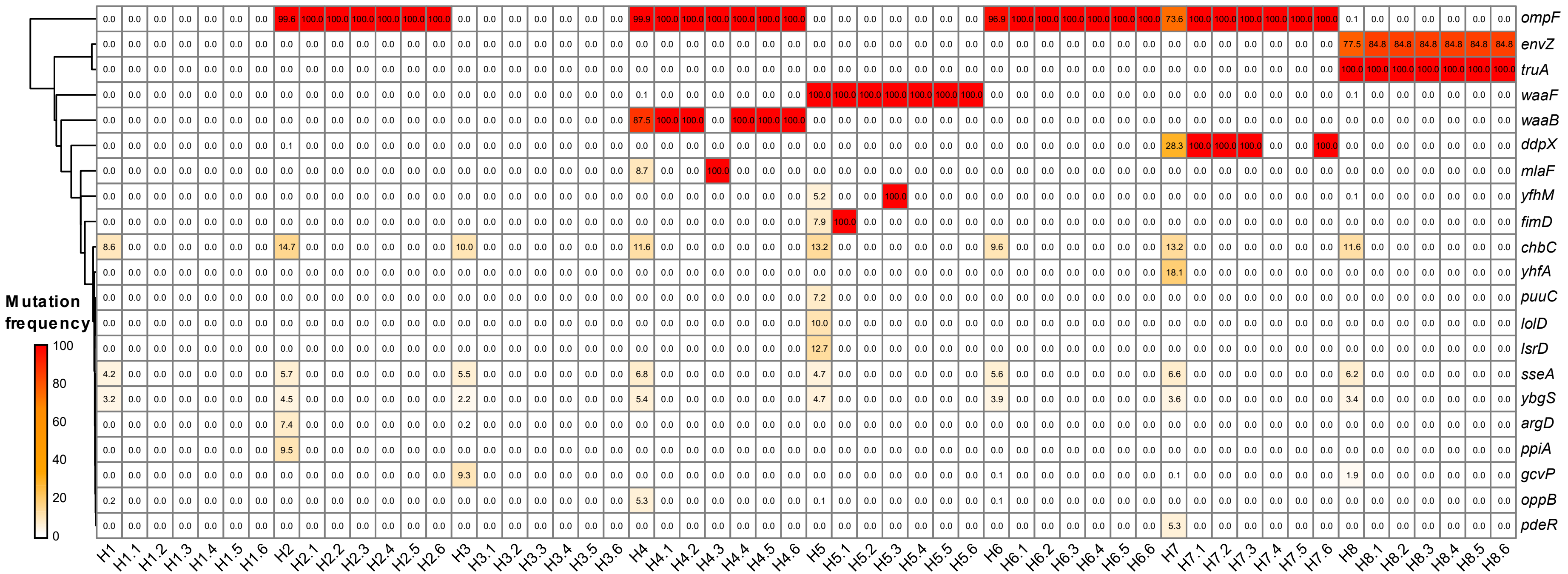


**Extended Data Fig. 9 │ Comparison of gene mutations between single clones and populations that evolved under high predatory pressure.** Hierarchically clustered heatmap of gene mutations with at least 5% frequency within any high predatory pressure evolved lineage populations H1-H8 (excluding rRNA and transposase genes, in Extended Data Fig. 7) at the 15$^{th}$ evolution cycle in comparison to the ancestor (AN). The 21 genes shown are exclusively mutated under high predatory pressure in H1-H8 populations. Next to each column of population sequencing data of H1-H8, is variant calling data shown from single clones 1-6 originating from that population (e.g. H1.1-H1.6). The population and single clone mutations are matched based on nucleotide position. For populations, mutation frequencies denote the actual percentage of a particular variant within a population. For single clones', the mutation frequencies shown are the highest mutation frequency of a particular variant, 0 indicates that variant is absent. The single clone sequencing data is analyzed with at least 3x coverage and a mutation frequency of at least 10%. All sequencing data was mapped against *E. coli* K-12 MG1655 prior to further analysis.

**Extended Data Table 1 │ Table of gene mutations in evolved prey populations.** Detailed characterisation of all mutations (>5% difference from the *E. coli* K-12 ancestor) within a coding gene in prey lineages H1-8, L1-N8) exclusively evolved under high and/or low predatory pressure (see Fig. 4). The same colour code was used as in Fig**.** 4 for the mutation type. Abbreviations: SNP: Single nucleotide polymorphism, InDel: Insertion or Deletion, fs: frame shift.

| gene | prey | % | impact | type | mutation type | change |
|---|---|---|---|---|---|---|
| *ompF* | H2 | 99.6 | moderate | SNP | missense variant | Ser294Pro |
| *ompF* | H4 | 99.9 | high | InDel | frameshift | Tyr332fs |
| *ompF* | H6 | 96.9 | high | InDel | frameshift | Glu70fs |
| *ompF* | H7 | 73.6 | high | InDel | frameshift | Asn183fs |
| *ompF* | H7 | 25.7 | high | InDel | frameshift | Tyr332fs |
| *ompF* | L1 | 00.0 | high | SNP | stop gained | Gln88* |
| *ompF* | L2 | 5.1 | high | SNP | stop gained | Gln88* |
| *waaF* | H5 | 100 | moderate | SNP | missense variant | Ala255Glu |
| *envZ* | H8 | 77.5 | moderate | InDel | conservative inframe insertion | Insertion of Lys-Thr-Trp-Leu between Leu135 and Lys136 |
| *truA* | H8 | 100 | moderate | SNP | missense variant | Val200Ala |
| *waaB* | H4 | 87.5 | high | SNP | stop gained | Glu17* |
| *ddpX* | H7 | 28.3 | low | SNP | synonymous variant | Val92Val (codon usage decreased) |
| *yhfA* | H7 | 18.1 | low | SNP | synonymous variant | Leu96Leu (codon usage decreased) |
| *sseA* | H1 | -6.2 | moderate | SNP | missense variant | Lys214Asn |
| *sseA* | H5 | -5.7 | moderate | SNP | missense variant | Lys214Asn |
| *sseA* | L2 | -5.2 | moderate | SNP | missense variant | Lys214Asn |
| *sseA* | L6 | -5.4 | moderate | SNP | missense variant | Lys214Asn |
| *yfhM* | H5 | 5.2 | low | SNP | synonymous variant | Val712Val (codon usage invariant) |
| *fimD* | H5 | 7.9 | low | SNP | synonymous variant | Pro838Pro (codon usage decreased) |
| *puuC* | H5 | 7.2 | moderate | SNP | missense variant | Asp388Glu |
| *lolD* | H5 | 10.0 | low | SNP | synonymous variant | His141His (codon usage optimised) |
| *lsrD* | H5 | 12.7 | low | SNP | synonymous variant | Gly187Gly (codon usage optimised) |
| *argD* | H2 | 7.4 | low | SNP | synonymous variant | Leu257Leu (codon usage decreased) |
| *ppiA* | H2 | 6.4 | high | InDel | frameshift | Thr85fs |
| *ppiA* | H2 | 9.5 | moderate | SNP | missense variant | Thr85Pro |
| *gcvP* | H3 | 9.3 | moderate | SNP | missense variant | Tyr109Ser |
| *ybgS* | H3 | -5.4 | moderate | SNP | missense variant | Thr121Pro |
| *mlaF* | H4 | 8.7 | high | SNP | stop gained | Gln175* |
| *oppB* | H4 | 5.3 | moderate | SNP | missense variant | Phe75Leu |
| *chbC* | H1 | -5.4 | moderate | SNP | missense variant | Phe60Cys |
| *chbC* | L4 | -5.3 | moderate | SNP | missense variant | Phe60Cys |
| *ypdE* | L4 | 8.0 | low | SNP | synonymous variant | Ala303Ala (codon usage optimised) |
| *ydjI* | L8 | 5.4 | moderate | SNP | missense variant | Asn139Lys |
| *yfbN* | L8 | 7.8 | moderate | SNP | missense variant | Ile18Ser |
| *yhjJ* | L3 | 5.9 | moderate | SNP | missense variant | Ile115Met |
| *yhjJ* | L3 | 5.2 | moderate | SNP | missense variant | Ile115Ser |
| *pdeR* | H7 | 5.3 | moderate | SNP | missense variant | Ala543Val |
| *yadL* | L1 | 5.6 | low | SNP | synonymous variant | Val166Val (codon usage optimised) |

#### **Additional online material:**

### Outer membrane changes enable evolutionary escape from bacterial predation

**Mridha *et al.***

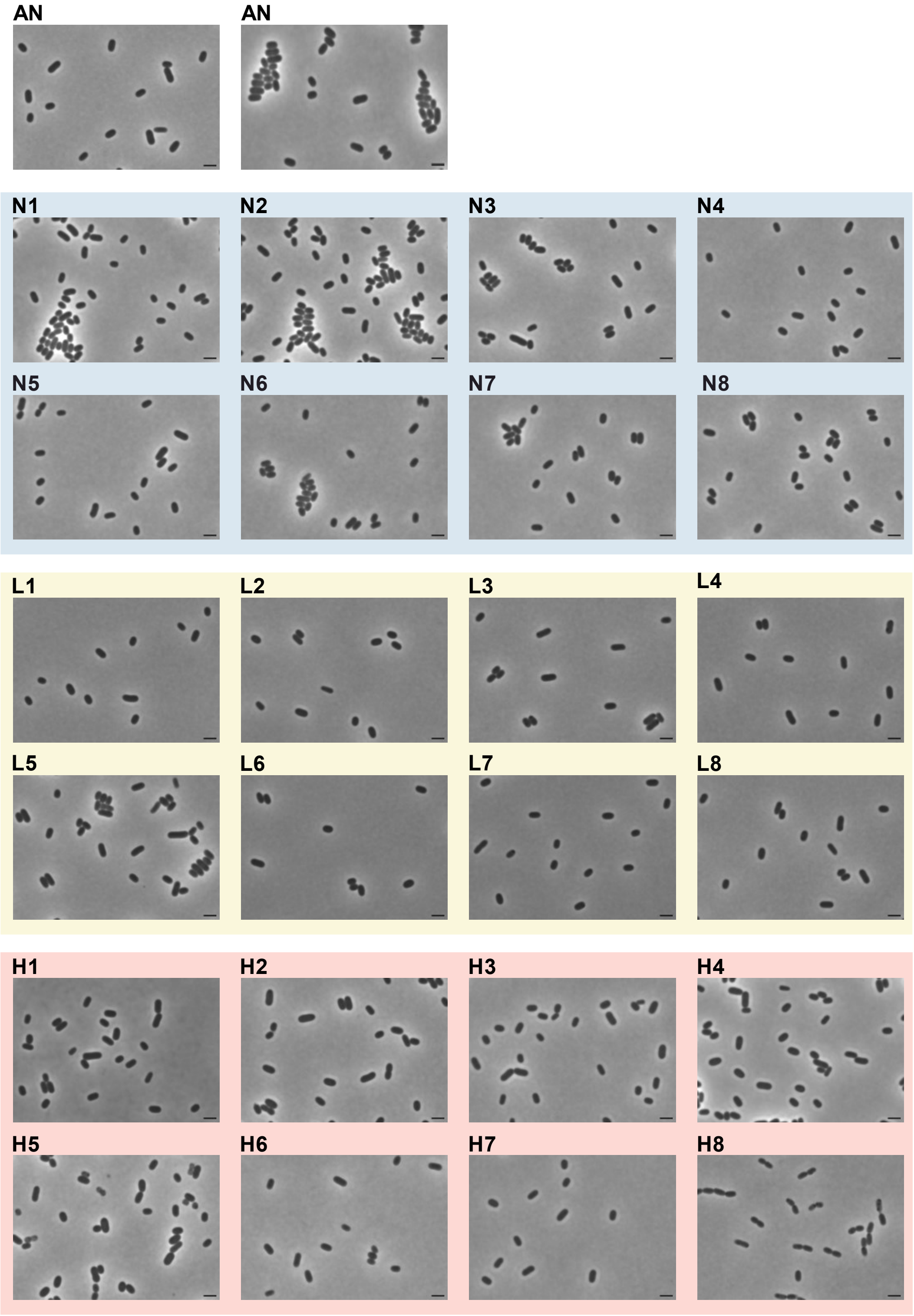


**Fig. SOnline1.** Phase contrast microscopy images of *E. coli* K-12 populations evolved under different predatory pressure for 15 evolution cycles show no morphological changes, except for line H8. Ancestor strain *E. coli* K-12 MG1655 (AN) and eight independently evolved lines under high (H1-8), low (L1-8) and no predatory pressure (N1-8) are shown. One representative image per evolved line was selected, while for AN two different ones are shown. One image of AN and H8 are also shown in Extended Data Fig S3. The scale bar is 2 μm.

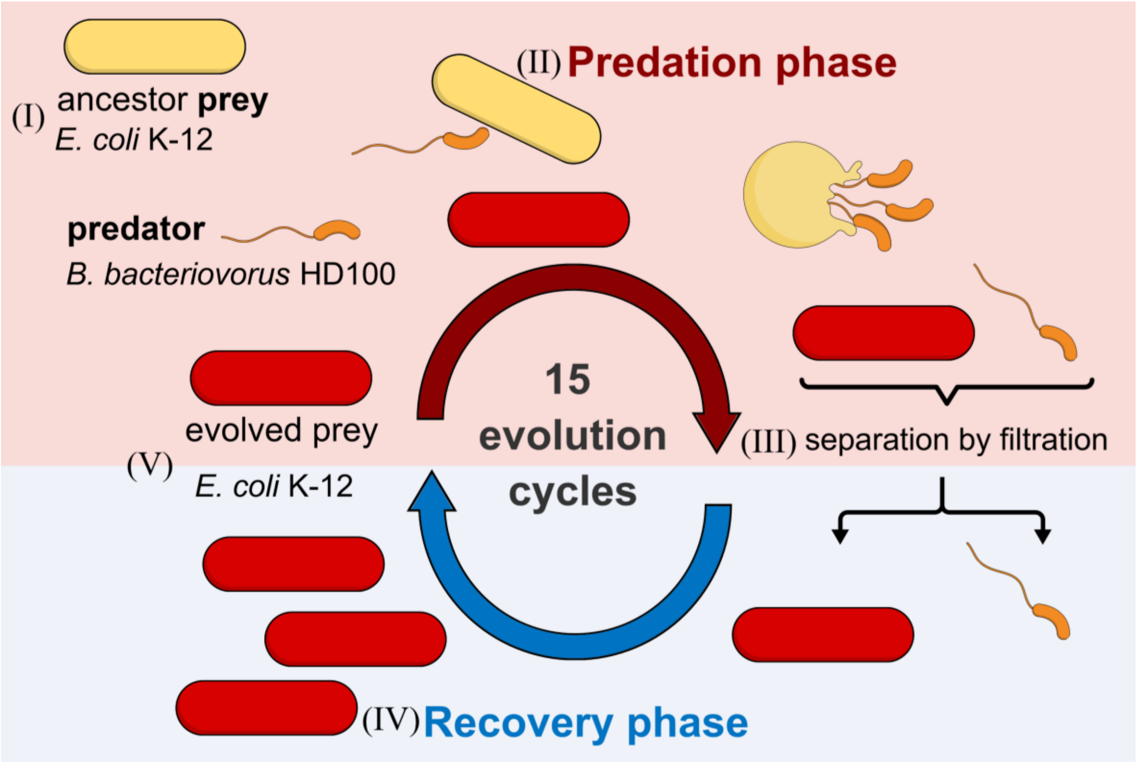


**Fig. SOnline2.** Overview experimental setup of prey evolution as shown in Figure 1a with added roman numbers for individual steps described at the end of this document.

**Table SOnline1.** *E. coli* strains used and generated in this study.

| **Strain** | **Genotype** | **Source** | **Reference** |
|---|---|---|---|
| *E. coli* K-12 MG1655 | wild type<br>$F^-$, $\lambda^-$, *rph*-1. Fnr+ | DSMZ (DSM 18039) | Blattner *et al.*[1] |
| *E. coli* K-12 BW25113 | wild type<br>*rrnB3, ΔlacZ4787, hsdR514, Δ(araBAD)567, Δ(rhaBAD)568, rph*-1 | Hall Lab, ETH Zurich | Baba *et al.*[2] |
| *E. coli* K-12 BW25113 *ompF*::*kanR* | Isogenic *ompF*::*kanR* | Hall Lab, ETHZ | Baba *et al.*[2] |
| *E. coli* K-12 BW25113 *waaF*::*kanR* | Isogenic *waaF*::*kanR* | Hall Lab, ETHZ | Baba *et al.*[2] |
| *E. coli* K-12 BW25113 *waaB*::*kanR* | Isogenic *waaB*::*kanR* | Hall Lab, ETHZ | Baba *et al.*[2] |
| *E. coli* K-12 BW2511 *envZ*::*kanR* | Isogenic *envZ*::*kanR* | Hall Lab, ETHZ | Baba *et al.*[2] |
| *E. coli* K-12 BW25113 *truA*::*kanR* | Isogenic *truA*::*kanR* | Hall Lab, ETHZ | Baba *et al.*[2] |
| *E. coli* K-12 BW25113 Δ*ompF* | Isogenic Δ*ompF* | This study | |
| *E. coli* K-12 BW25113 Δ*waaF* (*rfaF*) | Isogenic Δ*waaF* | This study | |
| *E. coli* K-12 BW25113 Δ*waaB* (*rfaB*) | Isogenic ΔwaaB | This study | |
| *E. coli* K-12 BW25113 Δ*envZ* | Isogenic Δ*envZ* | This study | |
| *E. coli* K-12 BW25113 Δ*truA* | Isogenic Δ*truA* | This study | |

**Table SOnline2**. Primers used to confirm clean deletion of *E.coli* BW25113 based on KEIO knock out library[2].

| strain | Primer name | Sequence (5' → 3') |
|---|---|---|
| Δ*ompF* | ompF_Seq_up | gaatggaaagatgcctgc |
| | ompF_Seq_dn | gttacagaagggaagtcc |
| Δ*waaF* | waaF_Seq_up | cttcaatctcggtactgg |
| | waaF_Seq_dn | ctgcgtcatagttctctg |
| Δ*waaB* | waaB_Seq_up | gctggactcttcgtatac |
| | waaB_Seq_dn | cataagcgatgtccagac |
| Δ*envZ* | envZ_Seq_up | cgtggtcgtgaatattcc |
| | envZ_Seq_dn | ctctgccgatgtttaacc |
| Δ*truA* | truA_Seq_up | ggtattctacggtcatgc |
| | truA_Seq_dn | taaatcaggtccatatttgc |

### Detailed protocols

##### Revival, cultivation of *Bdellovibrio bacteriovorus* HD100

We followed generally the protocol by Lambert & Sockett, 2008[3]. To revive *B. bacteriovorus* HD100 from frozen stocks, we used YPSC overlay plates. YPSC bottom agar plates are composed of 0.25 g $MgSO_4$ [Sigma], 0.5 g $CH_3COONa$ [Sigma], 1 g yeast extract [Bacto], 1 g peptone from meat, peptic digest [Sigma], 10 g agar [Sigma], per liter Mili-Q water, at pH 7.6, with added 0.25 g $l^{-1}$ $CaCl_2.2H_2O$ [Merck] after autoclaving. Then 5 ml of melted YPSC top agar (YPSC with 6 g $l^{-1}$ agar) at 55°C was mixed with 150 µl late-log *E. coli* K-12 MG1655 in a sterile 7-ml Bijou tube (Thermo Fisher Scientific) and poured on top of the YPSC bottom agar plate. After solidifying, 3 drops of 30 µl thawed predator stock from -80°C were placed onto the YPSC top agar and incubated at 29°C for 5 days. From the lytic zones of these YPSC overlay plates (indicative of *B. bacteriovorus* growth), material was transferred into 2 ml of Ca-HEPES buffer (5.94 g HEPES free acid [Sigma], 0. 294 g $CaCl_2.2H_2O$ [Merck], per liter Mili-Q water, buffered at pH 7.6) and supplemented with 150 µl of late-log *E. coli* K-12 MG1655 at $OD_{600nm}$ = 4.0 and incubated for 66 h at 29°C and 200 rpm. For the successive growth of predator, we transferred 50 µl of the previous lysate into 2 ml of Ca-HEPES buffer and 150 µl of overnight prey culture at $OD_{600nm}$ = 4.0. To ensure all *E. coli* prey cells were lysed after 66- or 24-h incubation in the lysate and free-swimming *B. bacteriovorus* had a fast-swimming speed suitable for fast predation, an aliquot was checked with a phase-contrast widefield microscope at 100x magnification.

#### Experimental prey evolution

##### *General setup*

In the experimental prey evolution, *E. coli* K-12 MG1655 was used as ancestral prey strain and subjected to three different predatory pressures (no, low, high) by *B. bacteriovorus* HD100 involving 15 alternate cycles of predation and recovery phase (Fig. 1a & Fig. Supplementary Online 2 with roman numbers referring to this text).

##### *I) Initial growth of E. coli K-12 MG1655 ancestor*

Prior to experimental evolution, we prepared overnight culture of *E. coli* MG1655 from a single colony. From this culture, we prepared 20% glycerol stocks and stored them in -80°C, to be used as reference for the ancestor strain (these ancestor glycerol stocks were used in the predation assays). To start the experimental prey evolution, we harvested 10 ml of this *E. coli* K-12 ancestor prey culture by centrifugation at 5311 x g for 3 minutes. The prey cells were resuspended in 0.8% NaCl and adjusted to $OD_{600nm}$ = 0.03. The prey and predator cultures were mixed in appropriate amounts in the subsequent predation phase.

##### *II) Predation phase*

A 24-hr lysate of *B. bacteriovorus* HD100 (described above in predator cultivation) was filtered through a 0.8-µm syringe filter (Whatman® Puradisc FP 30, Cellulose Acetate, sterile, WHA10462240) to remove prey debris and potential remaining prey cells. For the predation phase of the experimental evolution the cultivation media contained 30 ml YT broth, 10 ml 25 mM HEPES buffer and 2 mM CaCl2 (for the total volume of 40ml) enabling growth of *E. coli* prey as well as predation by *B. bacteriovorus*. Experimental evolution was performed in 24-well plates (Falcon, clear, sterile, non-tissue culture-treated, ref 351147). We used 3 different volumes of filtered predator lysate: 0 µl (no predator exposure), 40 µl (low predator exposure) and 160 µl (high predator exposure) and inoculated 150 µl of ancestor prey culture at OD600nm = 0.03, in 1.15 ml of cultivation media and adjusted the final volume to 1.5 ml per well, by either adding Ca-HEPES (as proxy for predator lysate) or 0.8% NaCl (as proxy for prey suspension). The final concentration of prey in reaction wells was OD600nm = 0.003.
The multiplicity of infection (ratio of predator (PFU/ml) vs prey cells (CFU/ml)) was ~ 2.5 for low and ~10 for high predatory pressure condition. For each predatory pressure condition, we had 8 replicates in total, distributed on two different 24-well plates, resulting in 8 independently evolving prey lineages for each experimental condition (no, low and high predatory pressure). As controls we made 4 replicates of each condition only without predator. We measured the

starting $OD_{600nm}$ of the reaction wells at the beginning of the predation phase in a multimode plate reader (Tecan Infinite M200 Pro), by shaking for 30 secs at 6 mm amplitude before measuring $OD_{600nm}$ with 10 flashes. In the predation phase the 24-well plates were incubated at 29°C and 200 rpm shaking (Infors Novotron AK85 shaking incubator) for 24 h. After the 24 h predation phase, we measured the end $OD_{600nm}$ of the reaction wells in the plate reader (same setting as before) and blank corrected the values against control wells containing only media (control wells without prey and predator). For each predatory phase a new 24-h lysate of *B. bacteriovorus* HD100 was used to avoid co-evolution, next to removing *B. bacteriovorus* at the end of the predation phase by filtration.

***III) Separation of surviving E. coli from B. bacteriovorus from predation phase by filtration***

At the end of each predation phase, we separated surviving prey from predator by filtration, using sterilized 0.45-µm 24-well filter plates (PALL, AcroPrep Supor, ref 97031) with a multi-well plate vacuum manifold (PALL, ref 5017). The filter plates were primed with 4 ml YT broth in each well for 15 minutes, then vacuum manifold set at pressure -10 in Hg for 1 minute was used to drain the YT broth. Based on the end $OD_{600nm}$ of the well after predation phase we transferred either 300 µl (when $OD_{600nm} > 0.30$) or 1 ml (when $OD_{600nm} < 0.30$) of the lysate from the predation phase plates onto the 0.45-µm 24-well filter plates and adjusted the final volume up to 1 ml with YT broth. The filtration was done at a vacuum pressure of -10 in Hg for 2 min. The residue (containing prey cells) was washed twice with 4 ml YT broth and filtered for 5 minutes, each at the same (previously mentioned) vacuum pressure. To facilitate filtration and avoid clogging of filter wells by prey debris we mixed each filter well by pipetting during the process. After filtration we resuspended the residue on top of the filter containing surviving prey cells in 1.5 ml YT broth and transferred it into new 24-well plates. We also measured $OD_{600nm}$ of the resuspended residue using the plate reader (same setting as before).

***IV) Recovery phase for E. coli prey***

For the recovery phase, we regrew the prey cells which survived the predation phase. We inoculated 15 µl of the resuspended residue into 24 well plates containing 1.5 ml of YT broth per well (~1:100 dilution). Again, we measured the starting $OD_{600nm}$ of the reaction wells (as previously). The 24-well plates were incubated at 37°C and 200 rpm for 16 h allowing surviving prey to grow in absence of the predator. To ensure no predatory pressure during this recovery phase (from a potential small amount of transferred predators) only YT broth, not containing any additional $CaCl_2$ was used. After the 16-h recovery phase, we measured the end $OD_{600nm}$ of the reaction wells in plate reader (same setting as before) and were blank corrected against control wells containing YT broth only. Then we prepared separate glycerol stocks of recovered prey containing 150 µl of evolved prey culture and 50 µl of 80% glycerol and stored them in -80°C in 96-well plates (Falcon, clear, non-tissue culture-treated, ref 351147).

***V) Starting new predation phase***

The recovered (evolved) prey was used to start the next evolution cycle, starting again with a predation phase as described for the first cycle using three different predatory pressures from a new 24-h lysate culture. After the predation phase, the surviving prey cells were again separated by filtering from the predator and the recovery phase followed as described for the first cycle.

We repeated the evolution cycles of predation and recovery phase 15 times overall. The predator used in every predation cycle, grown in Ca/HEPES at 29°C and 200rpm for 24 h, was always fed with 150 µl of $OD_{600nm} = 4.0$ overnight *E. coli* K-12 MG1655 ancestor strain. The cryo stocks of evolved prey lineages made after every recovery phase was later used in predation assays and for phenotypic and genetic comparison with the prey ancestor.

#### References:


1. Blattner, F. R., Plunkett, G., Bloch, C. A., Perna, N. T., Burland, V., Riley, M., Collado-Vides, J., Glasner, J. D., Rode, C. K., Mayhew, G. F., Gregor, J., Davis, N. W., Kirkpatrick, H. A., Goeden, M. A., Rose, D. J., Mau, B. & Shao, Y. The Complete Genome Sequence of *Escherichia coli* K-12. *Science* **277,** 1453–1462 (1997).

2. Baba, T., Ara, T., Hasegawa, M., Takai, Y., Okumura, Y., Baba, M., Datsenko, K. A., Tomita, M., Wanner, B. L. & Mori, H. Construction of *Escherichia coli* K-12 in-frame, single-gene knockout mutants: the Keio collection. *Molecular Systems Biology* **2,** 2006.0008 (2006).

3. Lambert, C. & Sockett, R. E. Laboratory maintenance of Bdellovibrio. *Curr Protoc Microbiol* **Chapter 7,** Unit 7B.2 (2008).
